# Lateral plate mesoderm directs human amnion and ventral skin organoid formation

**DOI:** 10.1101/2025.11.05.685987

**Authors:** Anh Phuong Le, Jin Kim, Qianyi Ma, Kelly Y. Gim, Sara A. Serdy, Edward H. Lee, Shariqa T. Shaila, Taiki Nakajima, Carl Nist-Lund, Yosuke Mai, Ian A. Glass, Laura C. Nuzzi, Catherine T. McNamara, Brian I. Labow, Liang Sun, Jiyoon Lee, Olivier Pourquié, Karl R. Koehler

**Affiliations:** Department of Otolaryngology; Plastics and Oral Surgery; F.M. Kirby Neurobiology Center, Boston Children’s Hospital, Boston, Massachusetts, 02115, USA; Department of Otolaryngology-Head and Neck Surgery; Surgery, Harvard Medical School, Boston, Massachusetts, 02115, USA; Program in Neuroscience, Harvard Medical School, Boston, Massachusetts, 02115, USA; Birth Defects Research Laboratory, University of Washington, Seattle, Washington, USA; Department of Genetics, Harvard Medical School, Boston, Massachusetts, 02115, USA; Department of Pathology, Brigham and Women’s Hospital, Boston, Massachusetts, 02115, USA; Research Informatics, Department of Information Technology, Boston Children’s Hospital, Boston, Massachusetts, 02115, USA; Center for Brain Convergence Research, Brain Science Institute, Korea Institute for Science and Technology (KIST), Seoul, Republic of Korea

**Keywords:** Organoids, Tissue Fabrication, Skin and Hair Biology, Amnion, Pluripotent Stem Cells, Developmental Patterning, Mechanobiology

## Abstract

Engineering organoids that faithfully replicate the intricate architecture and region-specific features of bodily organs and extraembryonic tissues remains a significant scientific challenge. Previously, we demonstrated that craniofacial skin organoids (cSkOs)—containing epidermis, dermis, and hair—could be generated by co-developing epidermal progenitors with cranial mesenchyme. Building on this approach, we precisely adjusted cellular composition and signaling environments to generate ventral skin organoids (vSkOs) with lateral plate mesoderm (LPM) progenitors, successfully recapitulating features of abdominal or groin skin. Modulating early BMP and FGF signaling redirected these vSkOs toward an extraembryonic fate, producing human amnion-like tissues, termed Amnioids. Like native human amnion, Amnioids rapidly expanded into large, avascular, hairless cysts, in sharp contrast to the primitive vasculature and abundant hair follicles of vSkOs. Single-cell RNA sequencing identified divergent molecular signatures and developmental trajectories, highlighting key roles for NOTCH, WNT, and YAP/Hippo signaling pathways. Functional studies further underscored mesenchymal-epithelial interactions and mechanical forces as critical regulators of epithelial expansion. Together, these models provide potent tools to investigate human development at the embryonic-extraembryonic interface, offering critical insights into congenital skin and amniotic disorders and opening new avenues for precision regenerative therapies.

## Introduction

During mammalian embryogenesis, the surface ectoderm (SE) initially forms a broad interface around the embryonic disk, giving rise to embryonic skin and the extraembryonic amnion—a fluid-filled sac protecting the embryo throughout gestation^1^. By week 4, rapid ventral body wall formation narrows this interface to the umbilical cord insertion as ventral skin expands to cover the abdomen, chest, limbs, and groin. Although skin and amnion share this common ectodermal origin, the developmental interplay between them remains poorly understood. Recent stem cell-derived embryo models have renewed interest in elucidating the molecular and mechanical pathways governing embryonic-extraembryonic ectoderm differentiation, with implications for developmental and regenerative biology^2–4^.

The diverse embryonic sources of the skin’s dermis are critical in determining region-specific characteristics^5–7^. Mammalian skin comprises an epidermis derived from SE and an underlying dermis arising from distinct embryonic lineages. For instance, the dermis of the face and scalp originates from cranial neural crest cells (CNCCs) and cranial mesoderm, respectively, yielding dense hair follicles and specialized glands^7–10^. In contrast, dorsal dermis arises from paraxial mesoderm, whereas ventral dermis covering the trunk and limbs derives from lateral plate mesoderm (LPM) (**Fig. 1a and Extended Data Fig. 1a**)^11–13^. Unique transcriptional signatures among these dermal populations suggest that dermal identity shapes epidermal differentiation and functional traits^8–10;^ however, this hypothesis remains untested in human stem cell-derived skin models.

**Figure 1:**
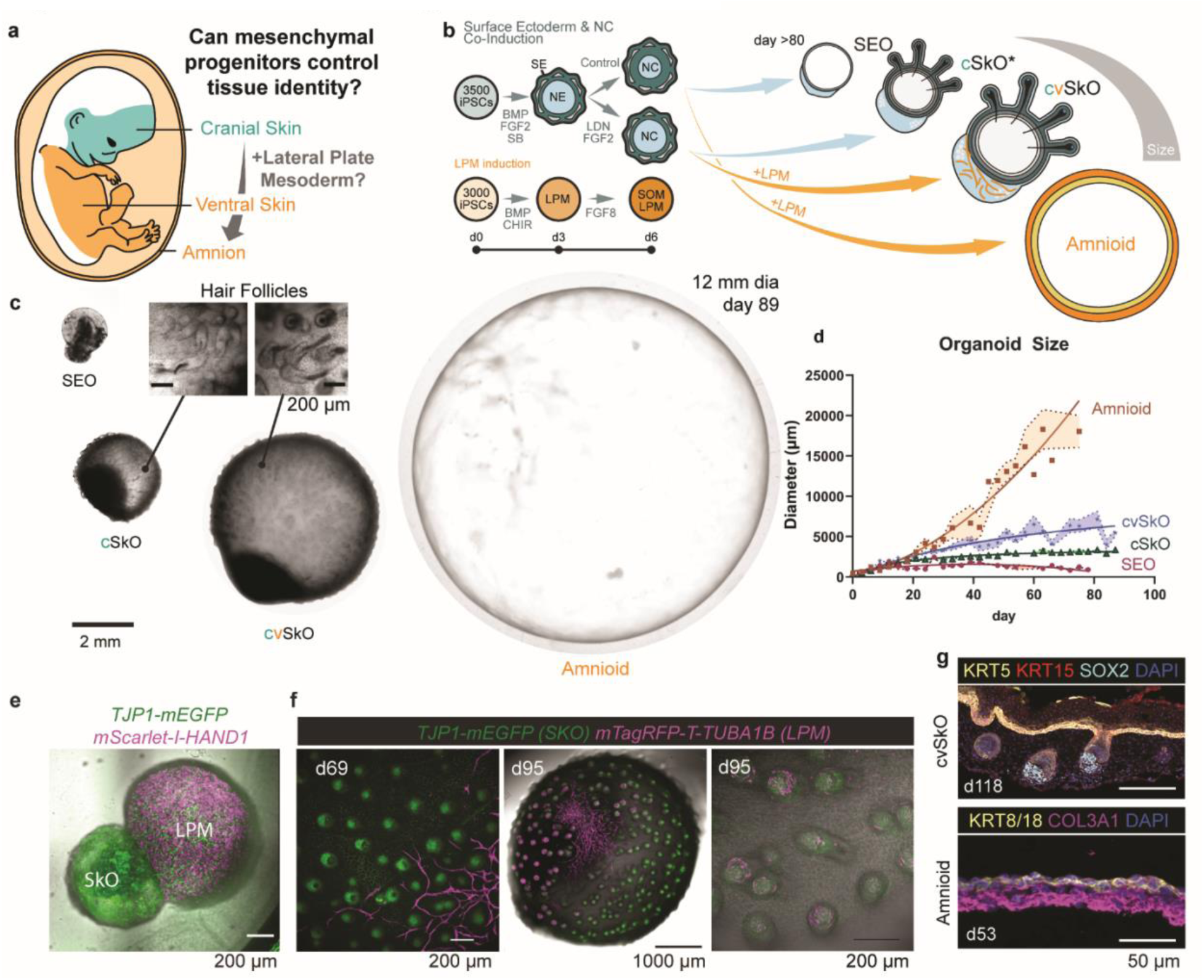
Generation of Trunk Skin Organoids and Amnioids. **a**. Schematic of developmental origins of craniofacial skin, ventral skin, and amnion, illustrating the hypothesis that mesenchymal cell identity influences epithelial fate. **b.** Experimental workflow for generating organoid models. Co-induction of surface ectoderm (SE) and neural crest (NC) yields cranial skin organoids (cSkOs), while inclusion of lateral plate mesoderm (LPM) generates ventral skin organoids (vSkOs) and Amnioids. NE: Neuroectoderm. SOM: Somatopleura Mesenchyme. **c.** Representative brightfield images of organoids at day 83–90 showing distinct morphologies. Skin organoids form hair follicles (insets), whereas amnioids develop large, follicle-free cystic structures (∼12 mm diameter). **d.** Quantification of organoid size over time demonstrates rapid expansion of amnioids relative to SEOs, cSkOs, and cvSkOs (SEO: n = 11 organoids, cSkO: 15 organoids, cvSkO: 24 organoids, amnioid: n = 13 organoids from 3 independent differentiation batches). **e.** Representative confocal image showing fusion of LPM (magenta) with SkO (green) to generate cvSkOs. **f.** Live confocal imaging from day 69 to 95 showing reporter expression in SkOs (TJP1-mEGFP, green) and LPM (mTagRFP-T-TUBA1B, magenta), revealing LPM integration and compartmentalization into vasculature, vasculature + dermal papilla, and dermal papilla regions (left to right). **g.** Immunostaining of organoid sections. Top: cvSkO showing epidermal markers KRT5 (green), KRT15 (red), and dermal papilla marker SOX2 (cyan). Bottom: day 53 Amnioid expressing amnion epithelial markers KRT8/18 (green) and amnion fibroblast marker COL3A1 (magenta). Both images are in the same scale.

Ventral skin has distinct features, including a larger surface area, delayed follicle initiation, and the absence of hair in specialized regions like palms, soles, and genitalia^14,15^. Interestingly, the LPM initially generates a continuous mesenchymal layer beneath the ventral skin and amniotic epithelium—known as the extraembryonic mesenchyme (ExE-Mech)—forming an avascular buffer between the amnion and chorion^16–18^. This anatomical proximity suggests ventral skin and amnion may share signaling and morphogenetic cues. Mechanical forces, increasingly recognized as regulators of skin morphogenesis, affect tissue expansion and hair follicle patterning^19–23^. Yet, how biomechanical factors shape mesenchymal lineage decisions and establish region-specific identities remains unclear. Given the early proximity between amnion and ventral skin, their mechanical environments might also influence their divergent morphologies.

Despite their developmental significance, ventral skin and amnion remain understudied in vitro due to a lack of models that replicate their distinct developmental dynamics. Existing craniofacial skin organoids (cSkOs) offer valuable insights but fail to recapitulate key features such as consistent vasculature and LPM-derived tissue characteristics^24–27^. To address this, we adapted our human pluripotent stem cell (hPSC)-derived cSkO protocol by incorporating LPM-derived cells and precisely controlling the initial cellular composition to generate ventral skin organoids (vSkOs) and amnion-like structures (Amnioids) (**Fig. 1b and Extended Data Fig. 1b**). This approach enabled investigation of developmental trajectories leading to ventral skin and amnion formation, highlighting the role of mechanical forces and signaling pathways, such as WNT, NOTCH and YAP/Hippo^22,23^. Our findings establish novel in vitro models to dissect embryonic-extraembryonic tissue interactions.

## Results

### Lateral plate mesoderm expands SkOs

In our previous study, the successful generation of cSkOs stemmed from the co-development of dermal and epidermal progenitors from CNCCs and SE cells, respectively^25^. Given the close spatiotemporal proximity of the LPM to ventral skin, we hypothesized that dermal progenitors differentiated from LPM cells could be incorporated into our cSkOs system to generate a new class of SkOs with more ventral trunk- or limb-skin characteristics (**Fig. 1a, b and Extended Data Fig. 1**). To validate this hypothesis, we utilized WTC-11 hiPSC lines (Allen Institute) carrying distinct fluorescent reporters to generate SkOs and LPMs in parallel. For LPM differentiation, we used a *HAND1-mScarlet-I/TJP1-mEGFP* double-reporter cell line, in which the fluorophore *mScarlet-I* was tagged into the *HAND1* gene to track lateral mesodermal progenitors (HAND1^+^/TJP1^-^), early epidermal progenitors (HAND1^-^/TJP1^+^), and SE or amniotic epithelia (AE, HAND1^+^/TJP1^+^). We induced LPM spheroids from hiPSCs through a combination of WNT and BMP4 activation (**Fig.1b & Extended Data Fig. 2a**), and subsequently directed their differentiation toward hindlimb mesenchyme – an LPM derivative that contributes to the ventral dermis of the posterior trunk and limbs – by treating with FGF8 (see **Supplementary Note 1** on LPM Derivation Strategy)^11,13,28–30^. To avoid anterior LPM derivatives, such as forelimb or cardiac mesenchyme, we omitted retinoic acid from the culture media (**Extended Data Fig. 2a-b**)^30–32^. Starting day 3 of LPM differentiation, we observed homogeneous HAND1^+^/TJP1^-^ cells within the LPM spheroid (**Extended Data Fig. 2b**). These cells also expressed BRACHYURY, confirming mesodermal identity, and showed downregulation of ECAD, consistent with complete epithelial-to-mesenchymal transition (EMT) from the pluripotent state (**Extended Data Fig. 2c**). We also confirmed the LPM lineage by the absence of TBX6 and KRT8/18 staining, indicating a lack of paraxial mesoderm (PXM) and SE, respectively (**Extended Data Fig. 2c**).

On day 6, LPM spheroids start to commit to a mesenchymal lineage by expressing PRRX1, a transcription factor associated with LPM specification and limb development (**Extended Data Fig. 2d**). Single-cell RNA sequencing (scRNA-seq) analysis of day 6 LPM revealed that our induced LPM-like cells exhibit a posterior LPM identity, characterized by strong *HOXA10*, *TWIST1*, *PRRX1*, and *HAND1* expression, with low *TBX4* (∼10%) and no *TBX5* or *TBX1* (**Extended Data Fig. 2e-g)**. The LPM gives rise to both body wall mesoderm, which contributes to the abdominal and chest dermis and limb mesenchyme, and *TCF21*+ *FOXF1+* visceral/splanchnic mesoderm, which forms mesenchyme associated with internal organs. Co-expression of *PRRX1* and *TWIST1*, together with low *TBX4*, absent *TBX5* and sparse *TCF21*, is consistent with a somatic LPM (body-wall)–biased state rather than a fully hindlimb-committed fate (**Extended Data Fig. 2e-f**). Additionally, 41.6% of LPM cells express high levels of *TBX3* and lack *FOXF1*, indicating that these cells are unlikely to be splanchnic mesoderm but may retain the potential to differentiate into extraembryonic mesoderm derivatives, such as the amniotic mesenchyme. A subset of 10.1% expressed markers consistent with hemangioblasts—such as *TAL1, ETV2, LMO2, KDR,* and *FLI1*—a specialized early endothelial cell population capable of generating *RUNX1*^+^ hematopoietic stem cells (**Extended Data Fig. 2e-f**)^30^. This subset of *RUNX1^+^* cells suggested endogenous hematopoietic potential within our LPM; however, we were unable to confirm this finding using immunostaining, likely due to their rarity. Nonetheless, TJP1-mEGFP^+^ endothelial cells became visible on the LPM spheroids between days 6 and 10 of differentiation (**Extended Data Fig. 2h**). By day 6, LPM spheroids contained HAND1^+^ mesenchyme with minimal epithelial cell presence, alongside posterior LPM features and a bias toward body wall mesoderm (**Extended Data Fig. 2a-b**). Given this transcriptional profile and developmental state, we selected day 6 as the merging timepoint with cSkOs generated in parallel (**Fig.1b, Extended Data Fig. 3a-b**) to both maximize co-developmental interaction time and align with a stage of high mesodermal plasticity.

We first asked whether incorporating LPM to our cSkO system could impart ventral-like features—such as increased size or delayed follicle initiation. We tracked the growth and hair follicle development of LPM-cSkO assembloids for more than 100 days. From day 7 to 12, LPM- derived mesenchymal cells migrated and enveloped the cSkOs (**Supplementary Video 1 & 2**). By day 80, the assembloids reached 6,966 ± 970.3 µm in diameter, approximately three times larger than cSkOs cultured without LPM, which averaged 2,114 ± 177.8 µm (**Fig. 1c-d & Extended Data Fig. 3c-d**). Unlike cSkOs, LPM-cSkO assembloids typically formed smaller mesenchymal tails with little to no cartilage-like tissue attached to the skin cysts (**Fig. 1c & Extended Data Fig. 3b & d**)^25,26^. Interestingly, hair germs emerged in the LPM-cSkO assembloids 7-10 days later than in cSkOs (**Fig. 1c, Extended Data Fig 3d-e**), mimicking delayed hair follicle initiation observed in ventral skin in vivo^33,34^. In addition, the LPM-cSkOs assembloids exhibited delayed epidermal stratification compared to cSkOs, characterized by thinner epidermal layers and discontinuous periderm at corresponding time points (**Extended Data Fig. 3f**). To assess LPM-derived cell contributions to skin development, we generated assembloids using the *mTagRFP-T-TUBA1B* hiPSC line for LPM and the *TJP1-mEGFP* lines for cSkOs (**Fig. 1e**), then monitored the mTagRFP-T fluorescent signal. The LPM-derived mesenchymal cells formed dermal fibroblasts, dermal papillae, and vasculature (**Fig.1f, Supplementary Video 3**). We also detected Peripherin (PRPH)^+^ sensory neurons (**Extended Data Fig. 3g**), indicating the presence of a significant population of cranial neural crest cells in the assembloids. Therefore, these LPM-cSkO assembloids have mixed cranial and ventral anatomical identities; we refer to them as cranial-ventral skin organoids (cvSkOs).

To further ventralize cvSkOs and confirm that LPM can fully generate the dermis, we attempted to eliminate the CNCCs by omitting LDN/bFGF treatment on day 3 of cSkO differentiation – a condition we previously showed to be critical for CNCC co-induction with SE cells in cSkOs^25^. The resulting SE organoids (SEOs), predominantly composed of SE epithelium (TJP1^+^/HAND1^+^) and lacking CNCC markers SOX10 and p75NTR (**Extended Data Fig. 4a-b**), did not mature into hair-bearing skin (**Extended Data Fig. 4c**) and reached an average diameter of 907 ± 266 µm by day 100 (**Fig. 1c-d**). To reconstitute a dermal component and test how LPM contributes as a mesenchymal cell source, we merged day 6 SEOs with day 6 LPM to generate LPM-SEO assembloids (**Fig. 2a** & **Extended Data Fig. 4d**). To our surprise, these assembloids expanded at an average rate of 230 µm/day, forming massive structures that reached an average diameter of 1.5 cm by day 60 (**Fig. 1c-d**, **Fig. 2a-b**). Throughout this period, they maintained a transparent unicellular epithelial membrane overlying 2–3 cell layers of mesenchymal cells (**Fig. 2a, Extended Data Fig. 4e**), which thickened to up to 100 µm by day 90 (**Extended Data Fig. 4f**). To support their continued growth, we progressively transferred these organoids to larger well plates (**Fig. 2a**), and eventually to 500 mL conical flasks (**Extended Data Fig. 4d**), where they expanded further to reach up to 3 cm in diameter (**Fig. 1c**). Notably, large-diameter LPM-SEO assembloids never formed hair follicles, even after 150+ days in culture, suggesting they did not acquire skin identity. We confirmed this phenomenon using a human embryonic stem cell line, WA25 (**Extended Data Fig. 4g-h**). Based on their unique characteristics, we hypothesize that LPM-SEOs represent the amniotic sac.

**Figure 2:**
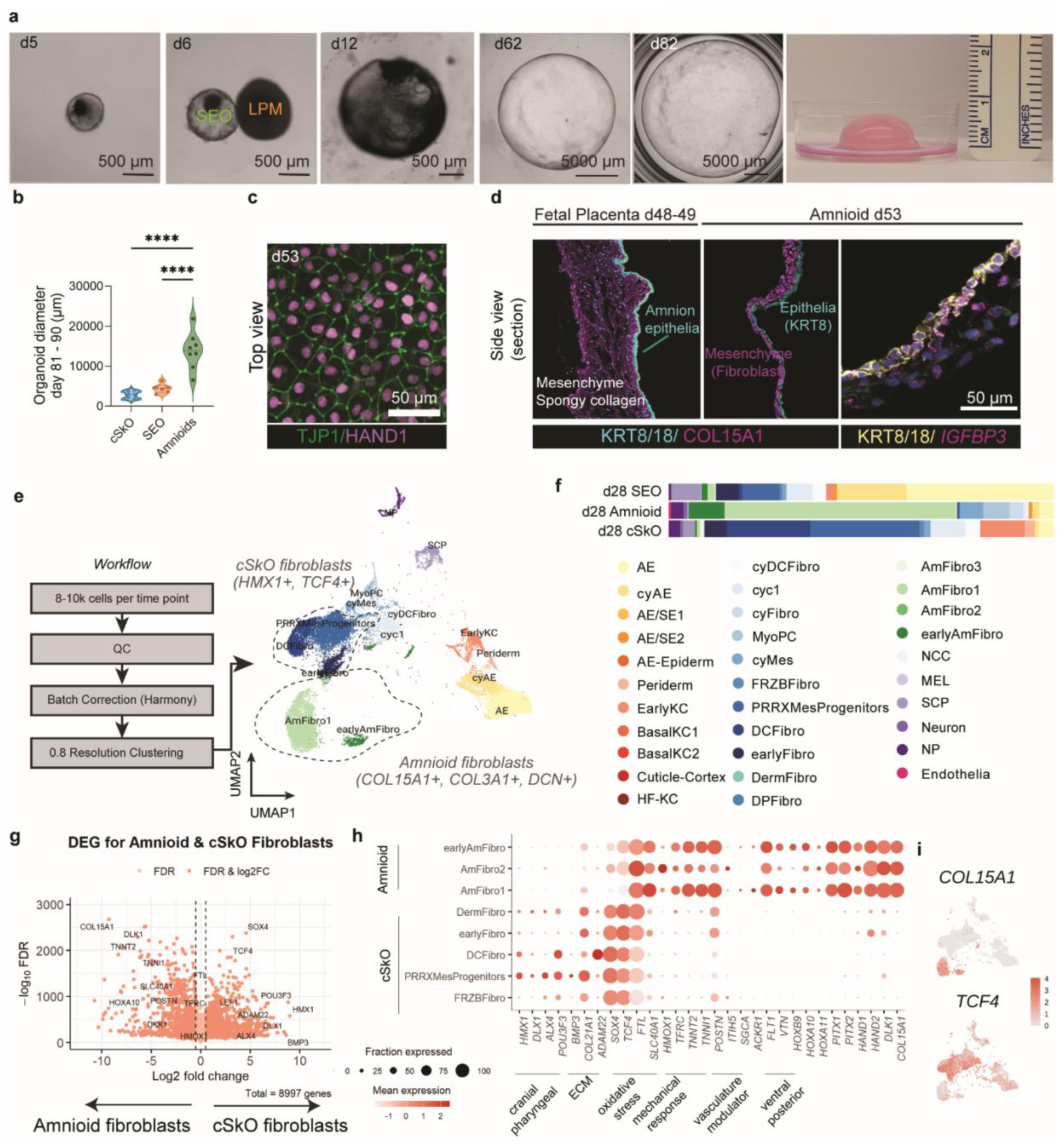
Differentiation and Characterization of Amnioids. **a**. Sequential bright field images showing Amnioid development following assembly of SEO and LPM on day 6. The last panel shows Amnioids in a petri dish, highlighting their macroscopic size. **b.** Quantitative size comparison showing Amnioids have largest size compared to LPM-free models (SEO and cSkO). (cSkOs: n = 10 organoids, 3 independent differentiation batches, SEO: n = 6 organoids, 3 independent differentiation batches, Amnioids: n = 8 organoids, 3 independent differentiation batches,) **c.** Confocal imaging of Amnioid epithelial layer (TJP1+) at day 53 post-differentiation showing HAND1+ expression, a marker of amnion identity. **d.** Validation of Amnioid identity through immunostaining and RNA in situ hybridization. Images show expression of KRT8/18 and *IGFBP3* (amnion epithelial markers) and COL15A1 (amnion fibroblast marker) in Amnioids compared with human fetal placenta. All images are in the same scale. **e.** UMAP visualization of single-cell RNA sequencing from Amnioids, SEO, and cSkO at day 28 post-differentiation, revealing distinct cell type populations across organoid models. **f.** Quantitative analysis of cell type proportions in each sample from panel e, showing model-specific cellular compositions. **g.** Volcano plot showing differentially expressed genes (DEGs) between Amnioid fibroblasts and cSkO fibroblasts (Venice test). **h.** DotPlot showing expression of top hits from DEGs analysis comparing Amnioid fibroblasts and cSkO fibroblasts. **i.** UMAP showing the differential expression of *COL15A1* as a distinctive Amnioid fibroblast marker, which is absent in *TCF4+* clusters (dermal fibroblast marker).

### LPM-SEO assembloids generate amnion

To clarify the identity of LPM-SEO assembloids, we compared their gene and protein expression profiles to those of native amniotic tissues. The epithelial layers consistently expressed early SE and AE markers, including KRT8/18 and HAND1, suggesting their commitment to an amnion-like fate (**Fig. 2c-d**)^35–37^. Supporting this notion, we observed prominent expression of *IGFBP3*, an established AE marker that is absent in cSkOs and human fetal skin samples (**Fig. 2d, Extended Data Fig. 5a**). These data strongly suggest the epithelial layer of LPM-SEO assembloids corresponds to authentic amnion. The presence of COL3A1, a known ExE-Mech marker associated with amniotic tissues, further supports the mesenchymal identity of amniotic origin (**Fig. 1g**). Additionally, sporadic expression of GATA4, a transcription factor associated with early ExE-Mech development, reinforces this interpretation (**Extended Data Fig. 4f**)^38^. Based on these epithelial and mesenchymal characteristics, we refer to LPM-SEO assembloids as “Amnioids.”

At day 28, scRNA-seq analysis revealed two distinct fibroblast populations with differential mechanical and signaling signatures (**Fig. 2e-f & Extended Data Fig. 5b-c**). Fibroblasts from cSkO expressed cranio-pharyngeal positional identity genes (*HMX1*, *DLX1*, *ALX4*, *POU3F3*), extracellular matrix (ECM) modulators (*BMP3*, *COL21A1, ADAM22*), and transcription factors promoting cell differentiation and hair folliculogenesis (*TCF4, SOX4*)^39,40^, indicative of a cranial dermal identity conducive to follicle formation (**Fig. 2g-h & Extended Data Fig. 5d**). In contrast, Amnioid fibroblasts highly expressed genes involved in mechanical tension, contraction, and stress responses (*TNNT2*, *TNNI1*, *POSTN*, *ITIH5*, *SGCA*), along with vascular ECM regulators (*ACKR1*, *FLT1*, *VTN*) and immature mesenchymal markers (*DLK1*). The presence of iron-handling and oxidative stress markers (*FTL, SLC40A1, HMOX1, CYGB*), together with posterior HOX factors (*HOXB9*, *HOXA11*), further highlights the specialized, developmentally distinct nature of the Amnioid stroma relative to dermal fibroblasts (**Fig. 2g-h, Extended Data Fig. 5e**).

Further analysis of the same day 28 scRNA-seq dataset–capturing an intermediate, expansion stage of Amnioids–identified unique fibroblast subpopulations distinct from those in cSkOs. Two transcriptionally defined clusters, earlyAmFibro and AmFibro1, both prominently expressed *COL15A1*–a marker not previously reported in ExE-Mech contexts–along with several transcription factors, *PITX1, PITX2, HAND1,* and *HAND2* (**Fig. 2h**). *COL15A1*^+^ cells also co-expressed known ExE-Mech markers *COL3A1* and *DCN*, while lacking expression of the dermal fibroblast marker *TCF4* (**Fig. 2h-i, Extended Data Fig. 5e**). Together, these molecular characteristics of Amnioid fibroblasts closely align with native ExE-Mech populations that contribute to the chorionic villi, connective stalk of the umbilical cord, and amniotic sac.

### Controlling size further ventralizes SkOs

We next sought to refine the generation of purer ventralized SkOs compared to cvSkOs by minimizing contaminating CNCCs, which may interfere with proper ventral identity acquisition; we refer to this new class of SkO as ventral SkOs (vSkO) hereafter. CNCCs arise from neuroectodermal cells located in the core of differentiating cSkOs. To limit this population, we reduced the initial size of aggregates, reasoning that smaller aggregates would allow cells to be more uniformly exposed to BMP4 signals, thereby promoting SE identity and reducing CNCC formation. As we decreased starting cell number from 3500 to 1600, 800, and 500 per aggregate, the resulting SkO cysts became more transparent (**Extended Data Fig. 6a**). By day 12, they exhibited comparable overall size **(Extended Data Fig. 6a-c),** but showed a lower mesenchymal-to-epithelial ratio (**Extended Data Fig. 6d**) and reduced numbers of SOX10^+^/p75NTR^+^ CNCCs (**Extended Data Fig. 6a**). Although aggregates from all starting size conditions could generate some hair follicle and neurons—indicative of CNCC contributions—these features were less frequent and consistent at smaller starting sizes (**Extended Data Fig. 6e**).

For subsequent experiments, we selected the 800-cell aggregates as the optimal starting condition, balancing minimal CNCC contribution with ease of handling. When combined with LPM on day 6, the resulting vSkOs developed into hair-bearing organoids of similar size to cvSkOs (**Fig. 3a-b**), exhibiting stratified epidermis (**Fig. 3c**) and comparable patterns of hair follicle formation (**Extended Data Fig. 6f-j)**. However, vSkOs contained fewer neurons (**Fig. 3d**) and fewer Schwann cells (**Extended Data Fig. 7a**), indicating a more effective reduction of CNCCs. Compared to cSkO, vSkO-derived hair follicles exhibited more uniform lengths and a blunter morphology, characterized by shorter length and a higher dermal papilla diameter-to-follicle length aspect ratio (**Fig. 3e–f, Supplementary Video 4-5**). To further confirm LPM contribution, we generated LPM using a *HIST1H2BJ-mEGFP* reporter line and SkO using an *mTagRFP-T-TUBA1B*. As *mTagRFP-T-TUBA1B* expression was suspected to diminish during differentiation, this dual-reporter approach enabled clear lineage demarcation (**Extended Data**ue characteristics, we hypoth). Broad distribution of mEGFP signals confirmed that LPM-derived cells populated the dermis and dermal papilla. Together, these data suggest that starting aggregate size influences mesenchymal composition and promotes complete contribution of the LPM to SkO dermis. Notably, Amnioids could also be generated from the 800-cell starting size (**Extended Data Fig. 7b-c**). Interestingly, 800-cell Amnioids showed few discernible differences compared to those initiated from 3500-cell aggregates (**Extended Data Fig. 7b-d**), suggesting that the Amnioid mesenchyme may suppress residual CNCC contributions, further supporting its developmental independence and distinct lineage identity.

**Figure 3:**
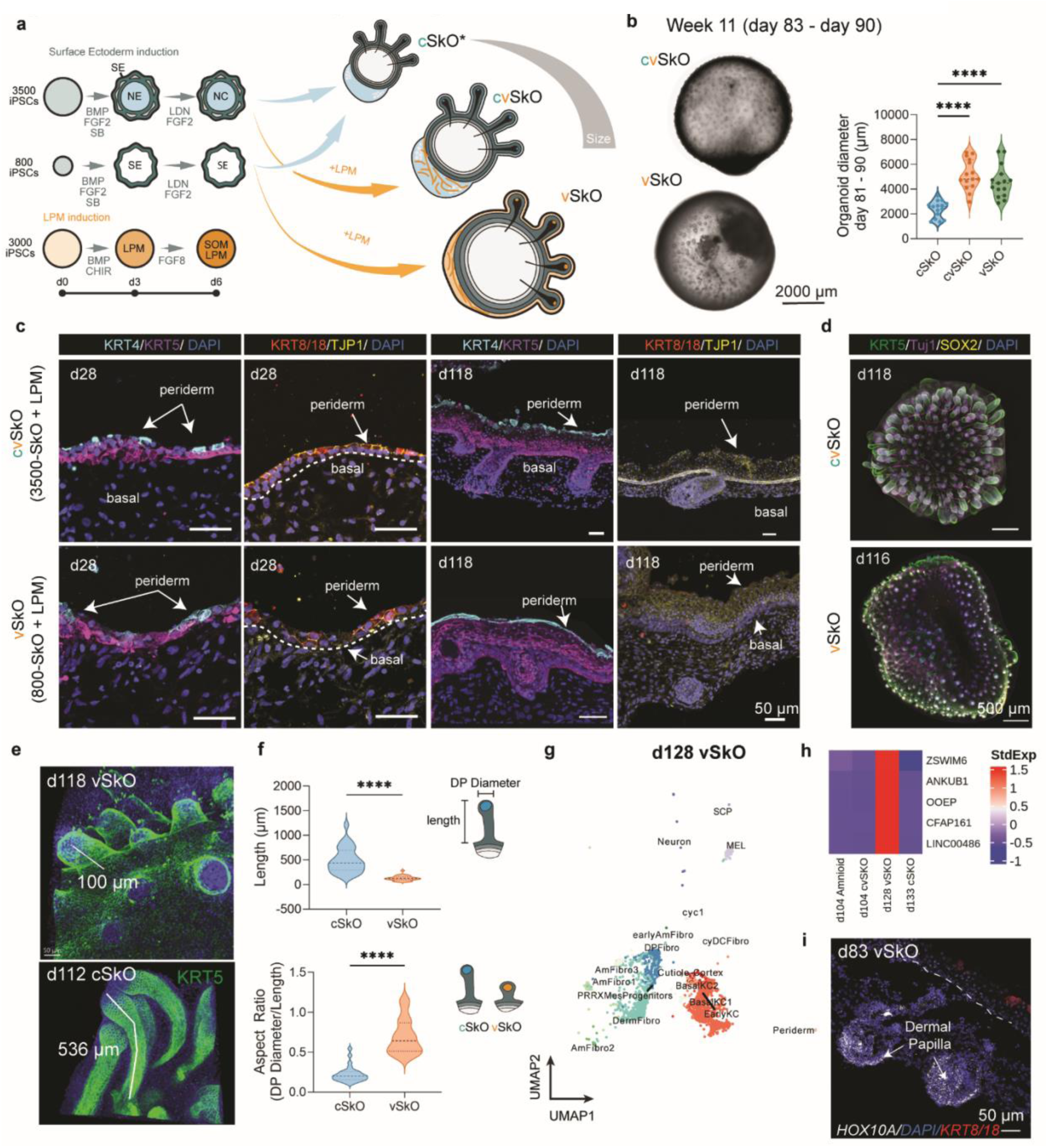
Ventralization of skin organoids by size control and assembly. **a.** Schematic of experimental strategy to generate vSkOs by reducing iPSC seeding density and assembling with LPM. **b.** Brightfield images of mature cvSkO and vSkO organoids at week 11 (days 83–90) show similar overall size and distinct surface morphologies. Violin plot quantification confirms comparable diameters (n = 15 organoids from 4 independent batches). **c.** Immunofluorescence of cvSkOs (top) and vSkOs (bottom) at early (day 28) and late (day 118) stages. Epithelial markers (KRT4/KRT5, KRT8/18) reveal proper stratification into periderm and basal layers. All images are in the same scale. **d.** Whole-mount immunofluorescence at day 118 (cSkO) and day 116 (vSkO) shows SOX2+ dermal papillae (DP) of hair follicles and reduced Tuj1+ neurons in vSkO. All images are in the same scale. **e.** 3D-rendered images of hair follicles from vSkO (top) and cSkO (bottom) at comparable stages illustrate shorter follicle length in vSkOs. **f.** Morphological comparison of hair follicles on day 112 – day 130. vSkO follicles are shorter and have wider DP, resulting in a stunted morphology. n = 28 cSkO follicles (5 organoids, 4 batches); 30 vSkO follicles (5 organoids, 4 batches). **g.** UMAP of scRNA-seq data from day 128 vSkO reveals distinct dermal and epidermal clusters. **h.** Heatmap of selected differentially expressed genes comparing vSkO and cSkO, highlighting ventral-specific transcriptional signatures in vSkOs. **i.** *HOX10A* expression detected by RNA in situ hybridization (RNAScope) in DP of day 83 vSkO, marking region-specific identity within early hair follicle-like structures.

To better define the anatomical location of vSkOs, we performed scRNA-seq on day 128 vSkOs (**Fig. 3g**), aiming to determine whether these organoids represent bona fide hindlimb skin or a neighboring posterior-ventral region, such as the abdomen or genital area. Notably, differential gene expression analysis of day 128 vSkOs (derived from male iPSCs) and day 133 cSkOs (from both female and male iPSCs) revealed a set of genes uniquely expressed in vSkOs. We noted several markers associated with testis development, such as *ZSWIM6*, *ANKUB1*, *OOEP*, and *CFAP161,* as well as the testis-associated long non-coding RNA, *LINC00486* (**Fig. 3h**). Importantly, RNAScope analysis of earlier-stage vSkOs (day 83) demonstrated *HOXA10* expression localized to dermal populations, indicating contribution from LPM-derived cells and specifying a posterior-ventral skin identity (**Fig. 3i**). Together, these findings suggest that vSkOs most closely reflect an abdominal or groin regional identity.

### Molecular signals direct Amnion vs. skin fate

Drawing from our single-cell analysis, we explored molecular signals controlling early fate acquisition and later maintenance in Amnioids and SkOs (cSkOs, cvSkOs, and vSkOs). Transcriptomic profiling at day 6 revealed that early BMP–FGF signaling establishes two distinct epithelial competence states (**Extended Data Fig. 7e**). Treatment with bFGF combined with BMP inhibition (LDN) activated transcriptional modules linked to epidermal stratification, ECM remodeling, and cell migration (*CD44*, *MMP9*, *BGN*)^41^, along with signaling integrators of Ca²⁺, cAMP, and ERK pathways (*TESC*, *PPP1R1B*, *KCNMB1*, *ADRB2*, *RCSD1*)^42,43^. Elevated expression of *TBX2*—typically expressed in lateral ectoderm^44,45^—suggested priming toward epidermis. In contrast, untreated epithelial precursors destined for Amnioids preferentially expressed canonical extraembryonic keratins (*KRT7*, *MFAP5*)^46^, endogenous BMP antagonists (*NOG*, *FST*)^47^, and genes involved in transport and barrier functions (*SLC15A2*, *SLC40A1*, *CLIC3*, *MUC15*)^47–49^. These cells also showed elevated expression of fluid-handling genes (*NPPB*, *CA2*), mechanical stress-response factors (*EGR1*, *ATF3*, *MAFF*, *IFI6*)^47,50,51^, and key transcriptional regulators of amniotic epithelium (*HAND1, GABRP, GATA3, SOX7*), aligning closely with recently defined amniotic signatures^47,52,53^ (**Extended Data Fig. 7f**). These profiles, visualized by heatmaps and UMAP overlays (**Extended Data Fig. 7e-g**), suggest that low BMP and FGF signaling conditions stabilize a nutrient-transporting, mechanically sensitive epithelial fate conducive to rapid cyst expansion, whereas exogenous FGF combined with BMP inhibition primes a posterior-ventral epidermal transcriptional state prepared for stratification, ECM remodeling, and eventual hair follicle induction upon assembly with LPM.

On days 28 (mid-stage differentiation) and 104-133 (late-stage differentiation), we further investigated mechanisms underlying the suppression of hair follicle formation and vascularization in Amnioids. We observed high expression of the secreted WNT inhibitor DKK1 in Amnioid mesenchyme compared to dermal fibroblasts in day 28 SkOs (**Fig. 4a-c**). DKK1 overexpression in developing mouse skin has been shown to completely abolish follicle initiation^54,55^. Consistently, *LEF1*, a key folliculogenesis regulator, was notably absent in Amnioids (**Fig. 4a-c**)^14,54,56^. Interestingly, at later stages, the inverse relationship between *DKK1* and *LEF1* expression was less pronounced (**Fig. 4d-e**), suggesting that these factors are more critical for follicle initiation rather than maintenance. CellChat analysis identified enrichment of *DLK1*, an inhibitory Notch ligand, particularly in cvSkOs and Amnioids (**Extended Data Fig. 8a-c**). DLK1 localized to amnion-like epithelia, transitional epithelial clusters, and LPM-derived fibroblasts—lineages absent or minimally present in cSkOs (**Fig. 4f**). Notably, canonical NOTCH signaling^57–59^ is essential for hair follicle induction, and DLK1 competitively inhibits this pathway. Immunohistochemistry validated stronger DLK1 expression in Amnioids, cvSkOs, and vSkOs compared to cSkOs, particularly in periderm and subepithelial fibroblast regions lacking follicles (**Fig. 4f**, **Extended Data Fig. 8d-e**). These findings suggest that LPM-derived mesenchymal cells suppress hair follicle formation by dampening local WNT and NOTCH signaling.

**Figure 4:**
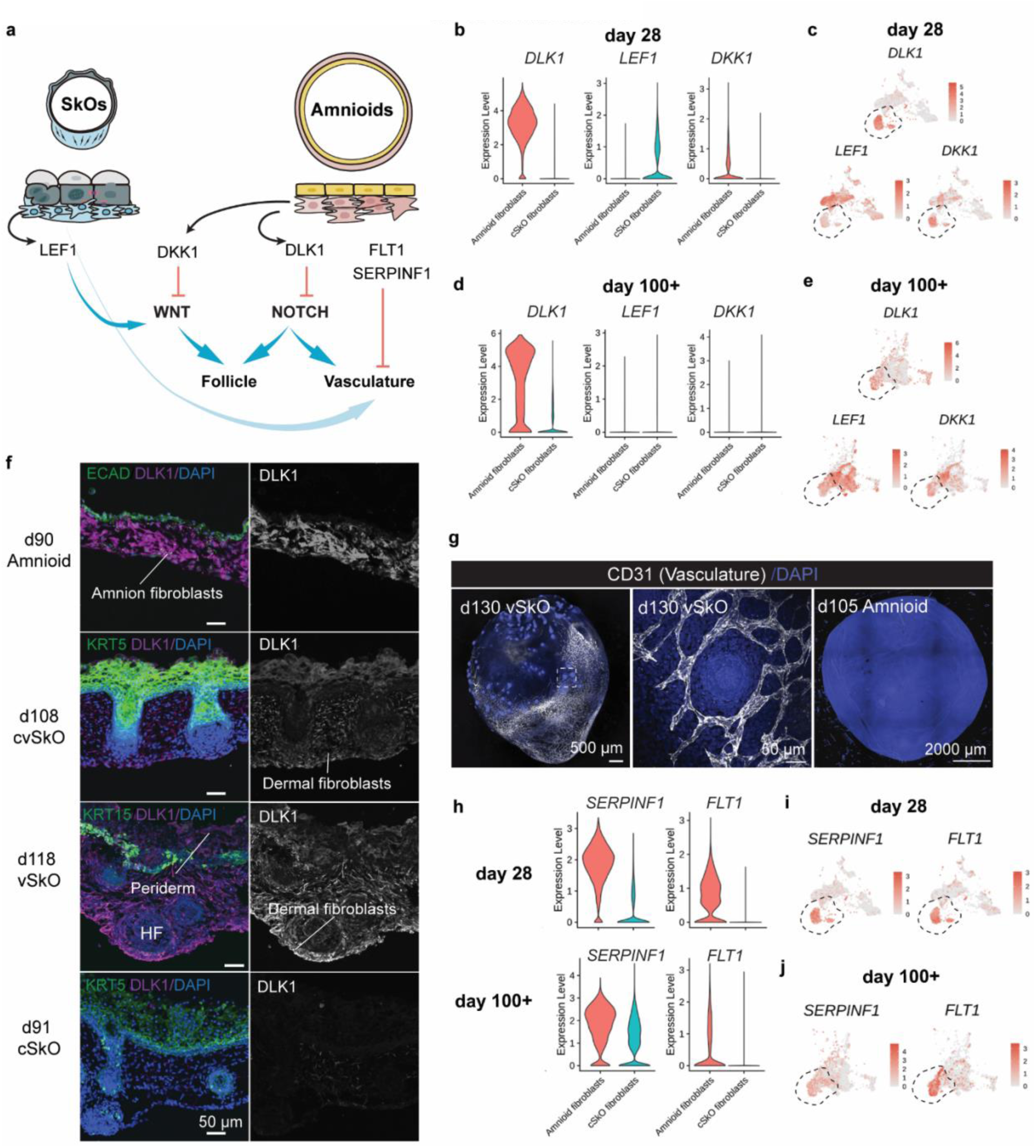
Mesenchymal subtypes control amnion vs skin fates. **a.** Schematic model showing how mesenchymal populations in SkOs versus Amnioids regulate epithelial fate via LEF1, DKK1, DLK1, FLT1, and SERPINF1. DLK1 suppresses folliculogenesis (via NOTCH), while FLT1/SERPINF1 is associated with vasculature formation. **b.** Violin plots of *DLK1, LEF1,* and *DKK1* expression on day 28 in single-cell data from Amnioids versus cSkO and vSkO conditions. **c.** Feature plots showing spatial expression of *DLK1, LEF1,* and *DKK1* at day 28. Amnioid fibroblasts clustered are shown within the black dotted line. **d.** Violin plots of *DLK1, LEF1,* and *DKK1* expression in 100+ single-cell transcriptomes. **e.** Feature plots showing *DLK1, LEF1*, and *DKK1* expression patterns at day 100+. Amnioid fibroblasts clustered are shown within the black dotted line. **f.** Immunofluorescence of DLK1 expression in fibroblasts from day 90 Amnioid, day 108 cvSkO, day 118 vSkO, and day 91 cSkO. DLK1 is present in Amnioid fibroblasts, dermal fibroblasts, and periderm. Scale bars, 50 µm. **g.** Immunostaining for CD31 in day 130 vSkO, cSkO, and day 105 Amnioid reveals limited vasculature in Amnioids. Middle panel shows magnified view of inset from left. Scale bars, 500 µm (left and middle), 2000 µm (right). **h.** Violin plots showing *SERPINF1* and *FLT1* expression at day 28 and day 100+ across epithelial-mesenchymal organoid types. **i.** Feature plots of *SERPINF1* and *FLT1* expression at day 28. Amnioid fibroblasts clustered are shown within the black dotted line. **j.** Feature plots of *SERPINF1* and *FLT1* expression at day 100+, enriched in mesenchyme of Amnioids and vasculature-associated populations. Amnioid fibroblasts clustered are shown within the black dotted line.

Regarding vascular development, LPM-infused SkOs exhibited robust vasculature, which was absent or minimal in Amnioids (**Fig. 4g**). Expression of *FLT1* (encoding soluble VEGFR1) and *SERPINF1* (encoding pigment epithelium-derived factor, PEDF)—potent anti-angiogenic factors known to maintain avascularity in tissues such as the cornea—was detected in the mesenchyme of Amnioids at days 28 and 104 (**Fig. 4h-j**). This suggests that a similar molecular mechanism actively preserves the avascular character of expanding Amnioid cysts, mirroring the naturally avascular human amnion.

To test whether modulating WNT signaling could shift early Amnioids toward a skin-like phenotype, we utilized our in-house tissue-chip platform (**Extended Data Fig. 8f-h**, which facilitates controlled co-culture and live-cell imaging of mesenchymal-epithelial progenitors. CHIR (WNT activator) treatment, initiated at day 18, induced mesenchymal condensation, reduced overall Amnioid size, and promoted epithelial thickening with enhanced keratinization (**Extended Data Fig. 8g-h**). This phenotypic shift indicated partial epidermal conversion. Immunostaining confirmed the emergence of stratified epidermis in the CHIR-treated day 18 Amnioids; however, hair follicle formation remained limited (**Extended Data Fig. 8g**). These results parallel previously reported WNT modulation phenotypes in mouse skin and underscore the interconnected regulation of amnion and skin specification^54^.

### Mechanical forces guide epithelial expansion

To explore how mesenchymal identity shapes epithelial architecture, we compared epithelial mechanics across Amnioids, SEO, cSkO, and cvSkOs. Each model exhibited distinct epithelial structures ranging from cohesive monolayers to stratified, follicular epithelia (**Fig. 5a**), suggesting differences in epithelial–mesenchymal mechanical coupling. Single-cell transcriptomics indicated that Amnioids and SEOs preferentially expressed programs related to organ growth, cell–cell junction assembly, and actomyosin contractility, whereas SkO models showed enrichment of EMT-related pathways (**Extended Data Fig. 9a**). Differentially expressed YAP-1 targets (*ANXA1– 3*), contractility regulators (*KANK4*), YAP inhibitors (*FRMD6*), and EMT markers (*CCN1, CXCL14, CD24*) further highlighted divergent mechanotransduction profiles.

**Figure 5:**
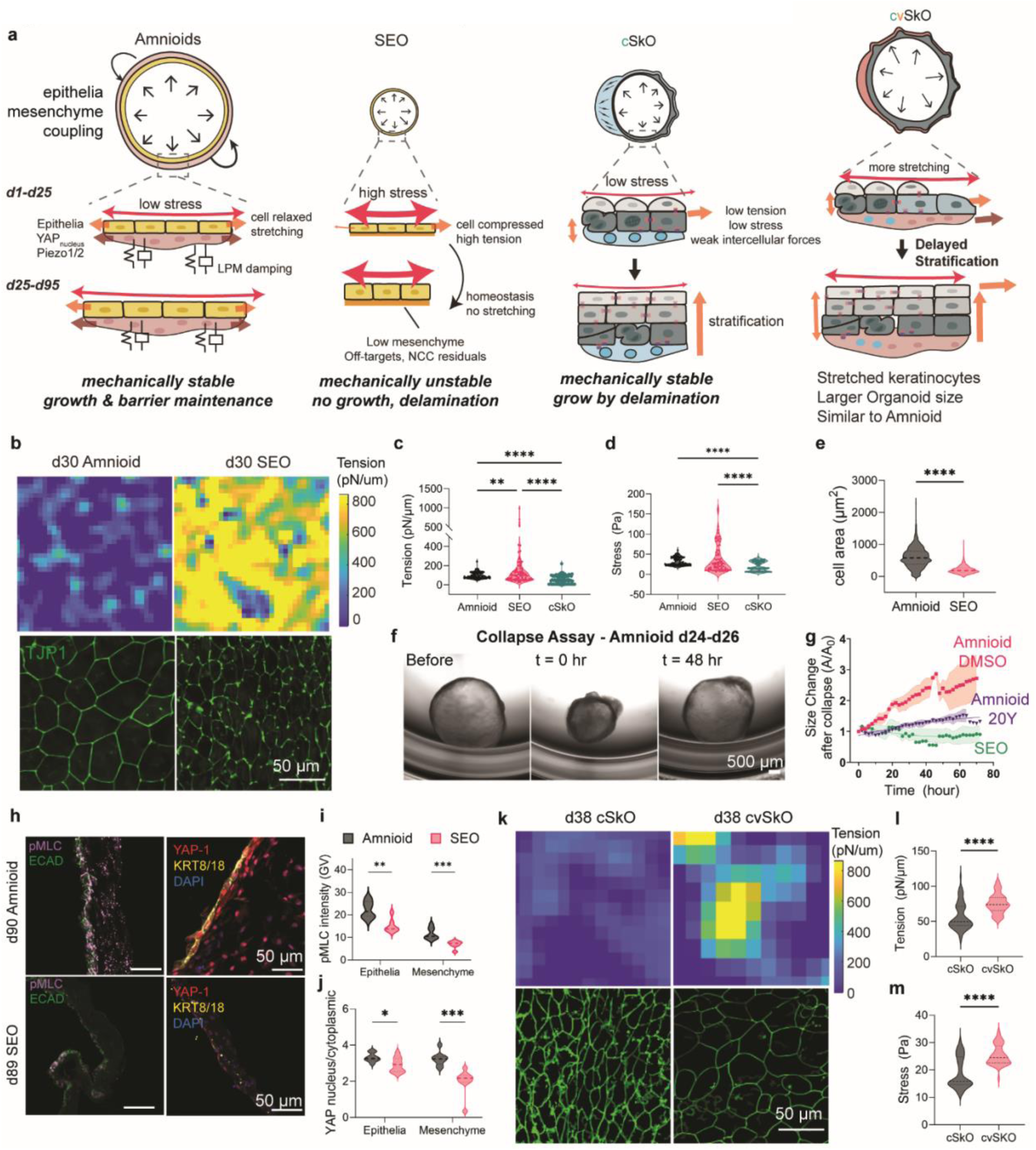
Mechanical stress dictates tissue expansion versus stratification. **a.** Schematic illustrating epithelial–mesenchymal mechanical coupling across Amnioids, SEOs, cSkOs, and cvSkOs. The presence of LPM-derived mesenchyme in cvSkOs renders them mechanically similar to Amnioids, promoting epithelial stretching and tissue expansion. **b.** Top: representative heatmap of tension measurements for day 30 Amnioids vs SEO. Bottom: representative images of tight junction protein signals (TJP1). SEOs are more mechanically unstable with higher tension overall and punctuated TJP1 at tricellular junctions. Scale bar, 50 µm. **c.** Quantification of tissue tension (pN/µm) in day 30 Amnioids, SEOs, and cSkOs. Each replicate is the average value from one timepoint of the time-lapse (Amnioids: *n* = 90 timepoints, 4 organoids, 2 across independent batches, SEO: n = 77 timepoints, 6 organoids, 4 independent batches, cSkOs: n = 104 timepoints, 4 organoids, 3 independent batches). **d.** Quantification of total mechanical stress (Pa) in the same conditions as in (c). **e.** Quantification of cell size (µm^2^) comparing between Amnioids and SEOs. Statistical comparison using unpaired student t-test (Amnioids: n = 1305 cells, SEO: 2960 cells from 4 organoids in 2 differentiation batches). **f.** Representative bright field image of day 24 Amnioid undergoing mechanical collapse and recovery. Scale bar, 200 µm. **g.** Time-course quantification of organoid size after being collapsed, comparing between amnioid, SEO and amnioid treated with 20 µM ROCK inhibitor Y27632 (20Y). **h.** Immunostaining for pMLC, ECAD, and YAP-1/KRT8 at day 90 Amnioids and day 90 SEOs. Amnioids show reduced actomyosin and nuclear YAP-1. Scale bars, 50 µm. **i.** Quantification of pMLC signals in epithelial and mesenchymal compartments (Amnioid: n = 8 sections, SEO: n = 7 sections from 3 organoids). **j.** Quantification of nuclear-to-cytoplasmic YAP-1 in epithelial and mesenchymal compartments (Amnioid: n = 8 sections, SEO: n = 9 sections from 3 organoids). **k.** Top: Tension heatmaps from day 38 cSkOs and cvSkOs. Bottom: TJP1 signal at the onset of stratification. Scale bar, 50 µm. The timepoint was chosen to match the start of stratification in SkOs. **l.** Quantification of tension (nN/µm) across day 38 cSkOs vs. day 38 cvSkOs (cSkO: n = 77 timepoints from 3 organoids in 2 independent differentiation batches, cvSkO: n = 52 timepoints from 3 organoids in 2 independent differentiation batches). **m.** Quantification of total stress (Pa) across day 38 cSkOs vs. day 38 cvSkOs (cSkO: n = 77 timepoints from 3 organoids in 2 independent differentiation batches, cvSkO: n = 52 timepoints from 3 organoids in 2 independent differentiation batches).

To link molecular differences to physical mechanics, we modeled epithelial layers as viscoelastic materials and quantified intercellular stress (σ) and tension (γ) and estimated these parameters via Particle Image Velocimetry (PIV) of the cell movements and cell morphology^60–62^ (**Extended Data Fig. 9b**). SEOs, which contain limited mesenchyme, exhibited the highest stress (59.32 ± 8.39 Pa) and tension (∼185 pN/µm), reflecting mechanically constrained, solid-like epithelia, whereas cSkO showed the lowest stress (22.62 ± 0.87 Pa), indicative of a mechanically relaxed state. Amnioids, which contain LPM, displayed intermediate mechanical values (σ = 31.58 ± 1.06 Pa) (**Fig. 5b–d, Supplementary Video 6**). These differences defined three distinct mechanical regimes of the epithelia: high-tension, solid-like SEOs; intermediate-tension, fluidic-like Amnioids; and low-tension, prone towards stratification cSkOs (**Extended Data Fig. 9c**). Supporting this, Amnioids exhibited significantly larger cell areas (2.3-fold, p < 0.001) (**Fig. 5b & e**), reduced packing density, increased phosphorylated myosin light chain (pMLC, 2.8-fold), and elevated nuclear YAP1 activity (1.9-fold) relative to SEOs, indicative of heightened mechanical plasticity and contractility (**Fig. 5h-j**).

Spatial correlation analysis in Amnioids revealed moderate-to-high correlations in tension (0.635 ± 0.037) and stress (0.656 ± 0.038) between the epithelial and mesenchymal layers, suggesting that the LPM-derived mesenchyme transmits epithelial forces while buffering epithelial motion **(Extended Data Fig. 9d-e, Supplementary Video 7)**. Mechanical perturbation assays further confirmed this buffering capacity: Amnioids recovered structural integrity within 48 hours after collapse, while SEOs did not re-expand **(Fig. 5f-g, Extended Data Fig. 10a, Supplementary Video 8)**. On-chip experiments showed that disrupting contractility (Y27632), adhesion (LLP-γ-PEG), or mechanotransduction (GsMTx4) impaired Amnioid expansion (**Extended Data Fig. 10b-c**), supporting mechanical control of epithelial architecture independent of lineage fate.

Introducing LPM into cSkO to create cvSkO resulted in elevated epithelial tension and stress (σ = 27.79 ± 0.81 Pa), reaching levels comparable to Amnioids (**Fig. 5k-m**). This intermediate mechanical regime correlated with enhanced epithelial expansion, delayed stratification, and postponed folliculogenesis relative to cSkOs (**Extended Data Fig. 3e-f & 6g-h**). Single-cell RNA-seq of day 104 cvSkOs revealed the retention of amniotic epithelial-like cells marked by persistent *HAND1* expression, along with elevated expression of mechanosensitive ion channels (PIEZO1/2) and YAP/Hippo targets^63^. These included genes involved in cell survival and proliferation (*ANXA1-3, WWTR1, TEAD1/4, BIRC5*), junctional and ECM stability (*AJUBA, TGM2),* cytoskeletal remodeling (*THBS1)*, and mechanosensing (*ANKRD1)* (**Extended Data Fig. 10d-e**). In summary, the delayed transition from epithelial expansion to stratification and follicle formation observed in cvSkOs and vSkOs—characteristic of trunk skin development—emerges from coordinated mechanical and molecular interactions between the LPM and the epithelium. These results underscore the critical role of mesodermal cells in orchestrating the mechanical and signaling environments required for proper tissue morphogenesis.

## Discussion

Our organoid models replicate key identities and functions of human amnion and ventral skin, offering new insights into how distinct epithelial fates emerge from shared precursors. By modulating BMP–FGF signaling and mesenchymal interactions, we show that LPM directs progenitors toward amniotic or epidermal fates. These findings validate longstanding yet largely untested hypotheses that mesenchymal lineage identity and mechanical signaling play critical roles in epithelial differentiation and tissue architecture. Our results are consistent with, and extend, recent studies highlighting mechanochemical feedback through YAP/Hippo and NOTCH signaling pathways, essential regulators of embryonic patterning and tissue morphogenesis.

While our vSkOs clearly exhibit characteristics consistent with abdominal skin, their exact anatomical identity remains incompletely defined. Future work employing enhancer mapping (e.g., ATAC-seq), single-cell epigenomics, and spatial transcriptomics (e.g., MERFISH) could precisely delineate positional identities within these organoids. Such analyses may uncover differential enhancer usage driving distinct gene-expression programs for LPM, amnion, and epidermis, deepening our understanding of early fate bifurcation and maintenance. Additionally, our single-cell analysis identifies *DLK1*—a known marker of early skin fibroblasts^56^ —as highly expressed in expanding Amnioid and vSkO fibroblasts, suggesting it may play a role in promoting expansion or inhibiting or delaying follicle formation. Given DLK1’s known role as a Notch inhibitor, it represents a candidate regulator of follicle development alongside known inhibitors, such as the WNT modulator DKK1. However, functional validation remains necessary.

The sensitivity of epithelial fate to mechanical perturbation emphasizes biomechanical regulation in our models. Mesenchymal cells serve as mechanical buffers, shaping epithelial expansion and stratification. Future organ-on-chip experiments varying ECM stiffness, adhesion, and mechanical forces could clarify how these parameters shape morphogenesis. Our models capture key developmental trajectories but have limitations. For example, their prolonged developmental timescales make it challenging to capture transient mechanical forces essential for epithelial morphogenesis. Our models also currently lack detail on ECM composition and basement membrane dynamics—key factors influencing mechanotransduction pathways. Addressing these gaps would further refine anatomical specificity to generate region-specific skin types, such as the fore and hindlimb skin or glabrous (hairless) skin of the palms, soles, and genitalia—areas rich in sensory nerve endings and challenging to regenerate^14,15,64,65^. Identifying mechanisms of rapid epithelial growth during fetal development could inform strategies to generate larger skin grafts for reconstructive surgery. It is intriguing to consider whether molecular and mechanical programs active in fetal amniotic expansion might be reactivated in adult skin during pregnancy or obesity, offering broader insights into tissue remodeling. In summary, our Amnioid and vSkO models enable detailed investigation of human development with therapeutic potential.

## Methods

### hPSC lines and culture

All culture experiments were performed with the WTC-11 hiPSC lines mTagRFP-T-TUBA1B, mEGFP-TJP1 (passages 35-55) acquired from the Allen Institute for Cell Science and the Coriell Institute. Cells were cultured on 6-well TC-treated plates coated with Vitronectin Recombinant Human Protein (Fisher Scientific) at a concentration of 0.5 µg/cm^2^. Pluripotent stem cells were maintained in mTeSR+ medium with supplement (STEMCELL Technologies) and 100 µg/mL Normocin (Invivogen; hereafter, mTeSR+). The medium was replenished every other day or everyday depending on cell confluency and passaged at ∼80% confluency (generally every 3-4 days). Cells were passaged in clusters of 3-6 cells on average with 0.5 mM EDTA.

### hPSC differentiation into cvSkO and vSkO

All steps take place in a biosafety cabinet unless stated otherwise. Pluripotent hPSC colonies were rinsed twice with 1x DPBS (Gibco), dissociated into single cells using StemPro Accutase Cell Dissociation Reagent (hereafter, Accutase; Gibco) and collected as a single cell suspension in mTeSR+ medium containing CEPT cocktail (final concentration in medium: 50 nM Chroman 1, MedChem Express; 5 µM Emricasan, SelleckChemicals; 1:1000 Polyamine Supplement, Sigma-Aldrich; 0.7 µM Trans-ISRIB, Tocris) and 100 µg/mL Normocin (Invivogen; here after: mTeSR + CEPT). On day minus 1, cells were distributed at a count of 3,500 cells in 100 µL mTeSR + CEPT per well for cvSkO and 800 cells in 100 µL mTeSR + CEPT per well for vSkO into 96-well U-bottom plates (Thermo Scientific). Aggregation was aided by centrifugation at RT, 110 rcf for 6 minutes. These cell aggregates were incubated in 37°C incubator under 5% CO_2_ for 24 hrs. On day 0, cells were transferred to new 96-well U-bottom plates in 100 µL of Essential 6 (Gibco) medium with 100 µg/mL Normocin (Invivogen; hereafter, E6), containing 2% Matrigel (Corning), 10 µM SB431542 (Reprocell), 4 ng/mL basic-FGF (PeproTech; herafter, bFGF), and 5 ng/mL BMP4 (PeproTech) to initiate non-neural ectoderm formation. On day 3, 200 µM LDN-193189 (BMP inhibitor, Reprocell; hereafter, LDN), and 50 µg/mL of bFGF in E6 medium were added in a volume of 25 µL per well, making the final concentration of 200 nM LDN and 50 ng/mL bFGF per well and the final volume 125 µL per well. On day 6 of differentiation, day 6 LPMs (generated in parallel) were merged with the SKO aggregates in 75 µL of E6 medium, making the final volume 200 µL. Half of the medium was changed (removal of 100 µL of spent medium and addition of 100 µL of fresh E6 medium) on days 8 and 10. On day 12, to induce self-assembly of epidermis, all aggregates were transferred into individual wells of 24-well low-attachment plates (Thermo Scientific) in 500 µL of organoid maturation medium (OMM) containing 1% Matrigel. OMM is composed of Advanced DMEM/F12 (Gibco) and Neurobasal (Gibco) media at a 1:1 ratio, 1X GlutaMAX™ supplement (Thermo Fisher Scientific), 0.5X B-27™ Minus Vitamin A supplement (Fisher Scientific), 0.5X N2 supplement (Fisher Scientific), 0.1 mM 2-Mercaptoethanol (Thermo Scientific), and 100 µg/mL Normocin (Invivogen). To maintain the aggregates in a floating culture for constant medium circulation, 24-well plates were placed on an orbital shaker shaking at a speed of 65 rpm in the 37°C incubator with 5% CO_2_. On differentiation day 15, half of the spent medium was replenished (removal of 250 µL of spent medium and addition of 250 µL of fresh medium) with OMM containing 1% Matrigel. Starting from day 18, half-medium change was performed every 2-3 days with fresh OMM without Matrigel, including once a week full-medium change. Increasing the total volume of OMM per well to 1-1.5 mL was necessary when the aggregates matured and grew larger to occupy approximately a quarter of the volume of the well. (See **Supplementary Notes** for additional differentiation protocol information and **Supplementary Table 1** for media compositions).

### hPSC differentiation into Lateral Plate Mesoderm (LPM)

On day –minus 1, colonies of hPSCs were dissociated and seeded onto 96 well U-bottom low attachment plates (S-Bio) at a density of 3000 cells in 100 µL per well with mTeSR + CEPT medium. Aggregation was aided by centrifugation at RT, 110 rcf for 6 min. On day 0, aggregates were individually collected and washed thrice with DMEM/F12 GlutaMAX™ (Thermo Fisher Scientific), 1% Insulin-Transferrin Selenium (Thermo Scientific; hereafter, ITS) and 100 µg/mL Normocin (Invivogen). Aggregates were then transferred to new 96-well U-bottom plates (S-Bio) in 150 µL medium of DMEM/F12 GlutaMAX™ medium containing 1% ITS, 100 µg/mL Normocin, 10 ng/mL BMP4 (PeproTech), and 3 µM CHIR99021 (Reprocell; hereafter CHIR). On day 3, medium was entirely replaced with 150 µL of DMEM/F12 GlutaMAX™ medium containing 1% ITS, 100 µg/mL Normocin and 10 ng/mL human FGF8a (PeproTech).

### hPSC differentiation into amnioids

On day –minus 1, colonies of hPSCs were dissociated similarly to SkO differentiation protocol on day –minus 1 and seeded at the density of 800 cells in 100 µL per well of mTeSR + CEPT medium. Aggregation was aided by centrifugation at RT, 110 rcf for 6 min. On day 0, all cell aggregates were individually collected, washed thrice with E6 medium and transferred to new 96-well U-bottom plates in 100 µL of E6 medium containing 2% Matrigel (Corning), 10 µM SB431542 (Reprocell), 4 ng/mL bFGF (PeproTech), 5 ng/mL BMP4 (PeproTech) to initiate non-neural ectoderm and amnion epithelia formation. On day 3, 25 µL of E6 medium was added to supply nutrients. For generating Amnioid (amnion epithelia + mesenchyme), on day 6, amnion epithelium was merged with day 6 LPM (generated in parallel) in 75 µL of E6 medium, making the total volume 200 µL. The plate was centrifuged at RT, 300 rcf for 5 min to assist the assembly of LPM with the amnion epithelia. From day 6 to day 9, fusion occurs by the migration of LPM cells to cover the surface of the amnion epitheila, making an assembloid with epithelia layer facing inside and mesenchymal cells outside. On day 8, half-medium change with E6 medium was carried out to supply more nutrients, and centrifugation can be performed if the organoids did not start to merge. On day 10, half medium was changed again. From day 12 onwards, the protocol was same as cvSKO and vSKO differentiation. To promote the growth, amnioids at day 30 to day 40 can be transferred to 6-well plates or bigger containers. Further notes on protocol variations and medium composition can be found in the Supplementary Information (page1-5).

### Mechanical collapse assay

Mechanical collapse assays were performed on organoids at day 24 post-differentiation, cultured in 24-well plates. To induce collapse, a single suction pulse was applied to the organoid membrane using a 200 µL pipette tip. This process was conducted under visual guidance using a benchtop microscope. Following collapse, organoids were immediately changed to 500 µL of fresh culture medium containing either DMSO (vehicle control) or 20 µM ROCK inhibitor Y27632 (Reprocell). Live imaging was initiated within 10 minutes of mechanical perturbation.

### Human Fetal and Adult Specimens

Facial skin tissue from an 18-week miscarried foetus was obtained from the University of Washington Birth Defects Research Laboratory. The tissue was not obtained from a living individual and was de-identified. De-identified adult human facial skin tissue samples were obtained from the Human Skin Disease Resource Centre at Brigham and Women’s Hospital-Harvard Medical School. Informed consent was obtained from all patients (or their legal guardians) who contributed samples used in this study. All specimens qualified for NIH Exemption 4 and were deemed "Non-Human Subjects Research" by the Indiana University School of Medicine IRB.

### Skin organoid dissociation for scRNA-seq

To dissociate organoids into single cells, randomly selected day 6 organoids (n = 24) and day 28 organoids (n = 10) were pooled from each group. For day 104 aggregates, two representative skin organoids were collected. Briefly, organoids before day 30 were incubated with pre-warmed (37°C) TrypLE™ (Fisher Scientific) for total 8-30 min (depending on the organoid sizes and densities) in a 37°C incubator on a shaker at 65 rpm for a gentle swirl. For day 104 organoids, organoids were first cut open and incubated with 10% Corning™ Dispase (Fisher Scientific) with CEPT for 30 min. During the incubation, the dissociation mixture was agitated every 3-10 min with gentle pipetting using wide-orifice p1000 and p200 tips (Mettler-Toledo) To aid the dissociation, organoids were pipetted at 10-minute intervals-first time gently with a wide-orifice p1000 (p200 for younger organoids; Rainin) tip and then with a regular p200 (Rainin) tip for subsequent intervals. For older organoids, if large cell chunks were remaining, the cell suspension was briefly spun down to collect larger chunks. The supernatant with Dispase-dissociated cells was saved in a separate tube while the larger chunks with treated with 300 µL of TrypLE™ with CEPT in the same manner as younger organoids. Once no visible chunks were remaining, the Dispase-dissociated cells suspension was mixed with the TrypLE™-dissociated cell suspension by pipetting. A neutralizing solution of cold 3% Bovine Serum Albumin (Millipore Sigma; hereafter, BSA) in culture medium with CEPT in a ratio of 1 (total cell suspension) : 2 (neutralizing solution) was added to the dissociated cell suspension to inactivate enzymatic activity. Then the cell suspension was centrifuged at RT, 230 rcf for 5.5 minutes, and resuspended in 1 mL of the cold neutralizing solution; this step was repeated twice. The suspension was then filtered through a 40 µm Flowmi™ cell strainer (Bel-Art™) into a 1.5 mL Eppendorf tube to eliminate any debris. Viability (live cell percentage) and live cell count was determined using Trypan Blue (Thermo Fisher Scientific), used at 1 (Trypan Blue) : 1 (Cell suspension) ratio, on a cell counting chamber slide and Invitrogen Countess II automated counter. Final cell concentration was ∼1,000 cells/µL and >90% viability.

### scRNA-seq cDNA library preparation and sequencing

Single cell 3’ RNA-seq experiments were conducted using the Chromium platform (10x Genomics) and the NovaSeq 6000 sequencer (Illumina, Inc). Approximately 17,000 cells (targeting for 10, 000 cells) per sample were added to a single cell master mix, following the Chromium Single Cell 3’ Reagent Kits v3 User Guide, CG000183 Rev C (10x Genomics). Day-6 samples were hash-tagged using CellPlex kit (10x Genomics) and pooled into one reaction of the Single Cell 3’ kit. Along with the single-cell gel beads and partitioning oil in separate wells of a Single Cell B Chip, the single cell reaction mixture was loaded to the Chromium Controller for GEM generation and barcoding, followed by cDNA synthesis and Illumina library preparation. At each step, the quality of cDNA and library was examined by Bioanalyzer. The resulting library was sequenced in on Illumina NovaSeq 6000 using 28b plus 91b paired-end sequencing to a read depth of ∼50,000 reads per cell.

### scRNA-seq data preprocessing

The raw single-cell RNA sequencing data was preprocessed using CellRanger v6.1.2 (10x Genomics), including aligning reads to the human reference genome (GRCh38-2020-A), and generating cellxgene expression count matrices. Using the generated expression count matrix/matrices, an in-house single-cell RNA-seq pipeline was built based on the Seurat R package (R v4.2.3, Seurat v4.3.0) (Butler et al 2018, Stuart et al 2019), including ambient RNA removal, quality control, cell filtering, spectral clustering, cell type annotation, differential gene expression, and visualization. Ambient RNA was removed using SoupX v1.6.2 (Young et al, 2020). Multiplets were identified using the scds v1.14.0 package using the cxds function (Bais et al, 2020). Only identified singlets were kept for further analysis. We filtered out cells exhibiting extremely low or high library sizes and number of gene features, falling outside the 95% confidence interval, as well as those displaying high mitochondrial content above the 10th percentile.

### Integration and clustering for entire data

Cells of good quality from the SE and cSKO samples at Day 6, 28, and 104 were merged and principal component analysis (PCA) over the identified 2000 highly variable genes was applied. Batch correction was conducted by integration using Harmony v0.1.1 (Korsunsky, Ilya, et al 2019). Cell clustering was performed on integrated data with a shared nearest-neighbor (SNN) graph-based method using the FindNeighbors function included in Seurat, followed by the Louvain algorithm for modularity optimization (resolution = 0.8) using FindClusters function. After the cell clusters were determined, their top 5 marker genes were identified with the FindMarkers function. For cluster annotation, the top marker genes based on the adjusted p value were manually curated to match canonical cell types and their marker genes based on literature research and public resources from scRNA-seq databases. The significance of cell type proportion differences was assessed using scProportionTest v0.0.0.9000 between all condition pairs. To perform the differential expression analysis, we utilized the Wilcoxon rank sum test with a logfc.threshold of 0.25 and a min.pct of 0.01, employing the FindMarkers function from Seurat. **Supplementary Tables 2-3** contain annotations and marker gene outputs)

### Integration and clustering at each developmental stage

Samples were separated into 3 groups based on developmental stages: 1) Early: d6 SE, d6 SE (+d3 LDN), d6 cSkO (SE + d3 bFGF/LDN), d6 LPM; 2) Mid: d28 SE, d28 Amnioid (SE + LPM d6 fusion), d28 cSKO (SE + d3 bFGF/LDN); and 3) Late: d104 Amnioid (SE + LPM d6 fusion), d104 cvSkO (cSkO + LPM d6 fusion), d128 vSkO and d133 cSkO. Samples at each of the 3 developmental stages were separately integrated using Harmony v0.1.1 (Korsunsky, Ilya, et al 2019) and clustering was performed as described above. For cell type annotation at each developmental stage, we first retained cell types present in the developmental stage based on a cell type proportion cutoff, which was determined as the intersect point of fitted Gaussian distributions to cell type proportions, and then performed label transfer from cell types present in the specific stage from the entire data using Symphony v0.1.1 (Kang, Nathan, et al 2021). The significance of cell type proportion differences was assessed using scProportionTest v0.0.0.9000 between all pairs at each developmental stage. **Supplementary Tables 4-5** contain outputs from differential gene expression analysis and NOTCH signal comparisons across samples and cell subtypes.

### Ion channel analysis

Ion channel gene list (N=341 ion channel genes in 44 ion channel families) was curated from the HGNC iron channel gene group (#177), IUPHAR/BPS Guide to pharmacology (IonChannelList), plus mechanosensitive ion channel family. Standardized log-transformed average expression level of top 10 differentially expressed ion channel genes in each cell type were visualized in heatmap.

### Gene set enrichment analysis

Differentially expressed genes between SE and cSKO in various cell types were enriched in the Gene Ontology (GO) database and the MSigDB Hallmark database using GSEApy v1.1.3 (Zhuqing Fang, Xinyuan Liu, Gary Peltz, 2022). The gp.gsea function was applied with 1,000 permutations to test for enrichment. Pathways with FDR q < 0.05 were considered significantly enriched. Enriched pathways were visualized using enrichment dot plots and bar charts.

### Cell-cell communication analysis

We used CellChat (version 2.1.2) along with its provided database of ligands and receptor pairs for humans to identify patterns of cell-to-cell communication between mesenchymal-epithelial cell types comparing SE and cSKO. To assess the likelihood of communication, we followed the methodologies outlined in the original research paper by Jin et al. (2024). We applied these methods at both the ligand-receptor pair level and the pathway level. To ensure reliable and significant findings, communication between cell types observed in fewer than 10 cells with adjusted p-values greater than 0.1 was excluded. The plots were generated using the built-in functions within CellChat.

### Flow Cytometry for LPM optimization

We used cell sorter to characterize LPM generated from WTC-11 *HAND1-mScarlet-I/TJP1-mEGFP* for the purpose of protocol optimization. 15 organoids were collected and enzymatically dissociated into single-cell suspensions using TrypLE Express (Thermo Fisher Scientific). Organoids were incubated in TrypLE at 37 °C for 15–20 minutes with intermittent pipetting every 3–5 minutes to facilitate dissociation. The resulting cell suspension was passed through a 40 μm cell strainer to remove undigested aggregates. Cells were resuspended in PBS supplemented with the CEPT cocktail (Y-27632, Chroman 1, Emricasan, and Polyamines) and kept on ice during transfer to the cell sorter to maintain viability and minimize cellular stress. Fluorescent cells were identified based on endogenous expression of mScarlet-I, without the need for additional staining. mScarlet-I⁺ cells were sorted using a BD FACSymphony™ S6 Cell Sorter (BD Biosciences) equipped with a 561 nm laser and appropriate filters. Gating strategy was implemented in BD FACSDiva v8.0.2 as follows: live single cells were first gated based on forward and side scatter (FSC-A vs. SSC-A), followed by doublet discrimination using FSC-H vs. FSC-A and SSC-H vs. SSC-W plots. mScarlet⁺ cells were identified in the PE-Texas Red-A channel and gated relative to a non-fluorescent control population (see **Supplementary Information – Supplementary Method Figure 1**). No antibody staining or viability dye was used. Sorted mScarlet⁺ cells were collected in complete culture medium on ice and processed within 1 hour for downstream applications.

### Immunohistochemistry

For immunostaining, fixed samples were cryoprotected through a graded treatment process of 15% and 30% of sucrose (Sigma-Aldrich) in 1x PBS, embedded on cryomolds (Endwin Scientific) in tissue freezing medium (General Data) or Optimal Cutting Temperature (OCT) compound (Fisher Healthcare), snap frozen and sliced to 12-15 µM thickness. For preparing sections of the Amnioids after day 24, the organoids were either cut open or collapsed and made into a Swiss roll using a thin wood dowel after sucrose gradient treatment. Cryosections were blocked in 10% normal goat/donkey serum (Vector Laboratories, hereafter NGS; Jackson Immunoresearch, hereafter NDS, incubated with primary antibodies diluted in 3% NGS/NDS, and then incubated with secondary antibodies in 3% NGS/NDS serum. Slides were mounted onto glass coverslip before imaging. Unless stated otherwise, images are representative of specimens obtained from at least 3 separate experiments. IHC analysis of later stages of development was performed on at least 3 aggregates from each condition per experiment.

### Wholemount immunostaining (SHIELD)

For wholemount immunostaining, samples were fixed in ice-cold 4% Paraformaldehyde (Electron Microscopy Sciences; hereafter, PFA) in 1x PBS for 12-18 hours and washed thrice with 1x PBS. The Clear+ Passive Clearing Kit (LifeCanvas Technologies) was used for reagents and buffers listed. Samples were then transferred on ice to 12.5% SHIELD-Epoxy-OFF solution at 4°C for 48 hours. The 12.5% SHIELD-Epoxy-OFF solution was prepared using 12.5 mL deionized water, 5 mL SHIELD buffer and 2.5 mL SHIELD-Epoxy solution. Then, samples were incubated in SHIELD-ON solution at 37°C for 24 hours. Following SHIELD fixation procedures, we then transferred organoids into a delipidation buffer (LifeCanvas Technologies) and incubated at 55°C for 24 hours. Once samples were ready for staining, samples were incubated with primary antibodies in 1x PBS with 0.1% Triton (Sigma-Aldrich; hereafter 0.1% PBS-T) for 18 hours. Following primary antibodies, samples were then incubated with secondary antibodies in 0.1% PBS-T for 18 hours. After antibody staining, the samples were incubated in graded Easy-Index matching medium (25%, 50%, 75% Easy Index in 1x PBS before 100% Easy Index) at 37°C for 18 hours.

### RNAScope

Samples were fixed with 4% PFA in 1X PBS. The tissues were then cryoprotected by sequential immersion in 10%, 20%, and 30% sucrose solutions in 1X PBS at 4°C until they sank. Samples were embedded in tissue freezing medium (General Data) or OCT (Fisher Healthcare) compound and frozen using dry ice or liquid nitrogen before storage at –80°C.

For sectioning, tissue blocks were equilibrated at –20°C in a cryostat and cut into 12–15 μm thick sections, which were mounted on SUPERFROST® PLUS slides. Sections were air-dried at –20°C for 60–120 minutes, washed with 1X PBS to remove OCT, baked at 60°C for 30 minutes, and post-fixed in 4% PFA for 1 hour at 4°C. Samples were dehydrated through graded ethanol (50%, 70%, 100%), air-dried, and processed for RNAscope® in situ hybridization following the manufacturer’s instructions, including target retrieval via steaming, probe hybridization, signal amplification, and fluorophore application. Slides were washed, counterstained with DAPI (Thermo Fisher), and mounted with Prolong Gold antifade (Invitrogen) for imaging.

### Organoid Morphometry Analysis

At least 10 organoids were pooled from at least three independent experiments and were used for recording organoid morphology over 100 days. From day 0 to day 12, every 3 days, bright-field images were taken by Nikon Eclipse Ts2R microscope using 4x objective to record the organoids morphology. From day 15 onwards, images of the organoids were recorded at least once per week to obtain the overall size and organoids morphology, using 2x objective on Nikon Eclipse Ts2R microscope or 0.5x or 1x objective on the Nikon SMZ18 stereomicroscope.

To quantify organoid size, the bright-field images were segmented using ilastik to obtain the overall outline of the organoids. The size was determined by the diameter of the fitted elipse of the outline. To quantify epithelial/mesenchymal proportion using bright field images, ilastik pixel classification was used to segment the epithelia (transparent parts) and the mesenchyme (dark, opaque parts).

### Hair Follicle Analysis in Bright Field Culture

*Frequency of organoids producing hair follicles (HFs).* At the end point of day 120 post-differentiation, organoids with protruding hair placodes, germs, pegs, and HFs were counted and recorded. Percentages of HF formation from at least 3 independent experiments per condition were calculated and averaged to get the general frequency of HF production from each cell line. *Timing of HF production.* At least three independent experiments were carried out to generate SKOs for tracking HF production. Bright-field images were taken at the frequency of at least once per week using benchtop microscope Nikon Eclipse Ts2R microscope at both 2x and 4x objectives. Among all organoids that produce hair follicles before day 120, images are traced back to the earliest day that hair germ was observed.

*Number of HF observed.* HFs number was estimated by counting based on bright-field images of organoids. As bright-field images can only show one side, organoids were checked thoroughly during imaging to obtain image of the side with most numbers of hair follicles.

### Hair follicles’ morphology analysis

Whole-mount 3D images of hair follicles (n=20) were acquired using 20x water-immersion objective imaging. Samples were labeled for keratinocytes (KRT5 or DSP), dermal papilla cells (SOX2 or HAND1), and nuclei (DAPI). 3D reconstructions were generated using Imaris v10.2. The Measurement Points tool was used to quantify hair follicle length and dermal papilla (DP) diameter. To measure follicle length, points were placed along the KRT5/DSP signal from the base of the follicle to the center of the dermal papilla. For DP diameter, two points were selected at opposite edges of the DP such that the line connecting them passed through its center, based on SOX2 or HAND1 signal. The Surface and ROI tools were used to assist with accurate point placement and structural delineation.

### Quantification of fluorescent signals

YAP1 and phosphorylated myosin light chain (pMLC) levels were quantified in paraformaldehyde-fixed tissue sections via immunofluorescence. Sections were co-stained with antibodies against YAP1 (Abcam, catalog #ab205270) or pMLC (Cell Signaling Technologies, catalog #3674S), along with E-cadherin (ECAD) to delineate epithelial regions and DAPI to mark nuclei. Nuclear YAP1 localization was assessed by segmenting nuclei using StarDist (ImageJ plugin), while cytoplasmic regions were defined by subtracting nuclear masks from the surrounding autofluorescence. Epithelial domains were segmented using the ECAD channel; mesenchymal regions were defined as the area remaining after excluding the epithelium from the total tissue area. Nuclear-to-cytoplasmic YAP1 ratios were calculated by dividing the mean nuclear fluorescence intensity by the mean cytoplasmic intensity. For pMLC quantification, mean signal intensities were measured separately in epithelial and mesenchymal compartments. All quantifications were performed on at least three independent tissue sections from biological replicates, imaged under identical settings using a Nikon A1R confocal microscope. Fluorescence channels and staining conditions were kept consistent across all samples.

### Quantification of cell size (segmentation)

To quantify cell size, we used organoids generated from *Hand1-mScarlet-I/TJP1-mEGFP* or *TJP1-mEGFP* cell lines to detect border by ZO-1 signals. Z-stacks of 4 µm of organoid images are taken to enhance ZO-1 cell-cell junction visibility during live imaging of organoids (within 12-18 hours), with 30 minutes interval. Images were then denoised using NIS Element Denoise.AI function, and post-processed to remove unspecific signals by training on ilastik pixel classification^66^ . Then, maximum projection. Maximum projected images are loaded onto Tissue Analyzer (PyTA: Python Tissue Analyzer v1.06a) ^67^ . Cell boundaries were detected using Deep Learning/EpySeg segmentation. Automated segmentation was followed by manual correction. Cell sizes were quantified using "Analysis" function, which provides measurements for individual cell area, aspect ratio, and elongation. Cell density, proliferation rate and T1 transitions were measured by a custom-made jupiter notebook.

### Quantification of cell dynamics and tissue mechanical properties

PIVLab v2.54, a Matlab plugin, was employed to analyze cell movements using the Particle Image Velocimetry (PIV) method. Z-stack images of 4 µm sections were first denoised, post-processed to eliminate non-specific signals, and corrected for 3D drifts. A maximum intensity projection was then applied to generate input data for PIVLab, where tissue-scale flows were tracked using a multi-pass windowed fast Fourier transform (FFT)-based approach. To minimize intensity variations and reduce tracking errors, contrast-limited adaptive histogram equalization (CLAHE) was performed with a 64-pixel box size. The PIV analysis was configured with an initial pass using a 64-pixel interrogation area with 50% overlap, followed by a second pass with a 32-pixel interrogation area and 50% overlap. Velocimetry data, including speed, correlation length, and direction, were analyzed using customized Matlab scripts.

Epithelial tissue stress analysis was conducted using a Maxwell viscoelastic model on PIV data, where strain tensor components (ɛ_xx_, ɛ_yy_, ɛ_xy_) were extracted from displacement fields and filtered before applying the Maxwell model. The constitutive equation, Dσ/Dt+σ/η=E⋅Dɛ/Dt, was solved using forward Euler integration with time steps matching image acquisition intervals to derive the stress tensor values (σ_xx_, σ_yy_, σ_xy_). Tissue-specific viscoelastic parameters were estimated using a Bayesian framework, with priors set for Young’s modulus (E) ranging from 100 to 5000 Pa (mean: 500 Pa, std: 300), Poisson’s ratio (ν) from 0.46 to 0.499 to reflect near-incompressibility, and viscosity (η) from 10 to 5000 Pa·s (mean: 500 Pa, std: 150). These priors, based on previous studies on embryos, were further refined using cellular morphological metrics measured by Tissue Analyzer. Labeled cells were analyzed to extract morphological parameters, including area, perimeter, roundness, aspect ratio, and eccentricity, which informed the Bayesian inference of mechanical properties: the area variation coefficient adjusted E to account for mechanical heterogeneity, mean cellular eccentricity modulated ν to capture shape-dependent compressibility, and cell packing density scaled η to represent emergent viscoelasticity. Theoretical strain responses were computed, and residuals between experimental and modeled strains were used in likelihood estimation, incorporating cellular penalties to improve inference accuracy. Markov Chain Monte Carlo (MCMC) sampling was performed with 10,000 iterations, including a 5,000-sample burn-in phase, to generate posterior distributions of E, ν, and η. Convergence diagnostics, including the Gelman-Rubin R_hat statistic, were computed to ensure robust parameter estimation. From the computed stress tensor components (σ_xx_, σ_yy_, σ_xy_), surface tension was derived as (σ_xx_ + σ_yy_)/2 * tissue thickness (converted to pN/μm). Statistical analyses included spatial and temporal averaging with standard deviations, with stress and tension distributions visualized as heatmaps. This framework integrates tissue-scale mechanics with cellular architecture, providing quantitative insights into the forces driving epithelial morphogenesis.

### Statistics and Reproducibility

Sample size was not predetermined using statistical tests, and investigators were not blinded to treatment groups. Experiments were generally not randomized; however, organoid specimens for histological and single-cell RNA sequencing analyses were randomly selected without prescreening for quality or morphological characteristics. Organoids used for measurements were derived from at least two independent differentiation batches. Statistical analyses were performed using MATLAB and GraphPad Prism 8. Data are presented as mean ± SEM unless stated otherwise. Violin plot limits represent maxima and minima, with individual data points shown as dots. If n > 30, individual data points are not shown on violin plots. Cell morphology metrics were pooled from at least three images per condition across three independent biological replicates (three organoids). Mechanical property and cell dynamics measurements were averaged per time point and pooled from three independent biological replicates. Comparisons between conditions were performed using unpaired 2-tail t-tests. Multiple comparisons were performed using Brown-Forsythe and Welch ANOVA with Durnett T3 pair-wise comparison.

### Use of LLM for Manuscript Preparation

The preparation of this manuscript involved the use of a Large Language Models (LLM), specifically OpenAI’s chatGPT-4.5, to assist in editing and refining the text where necessary. The scientific content, including the design of experiments, interpretation of results, and discussion of findings, was provided and thoroughly reviewed by the authors. All final decisions regarding the content and phrasing of the manuscript were made by the authors, ensuring the accuracy and integrity of the scientific information presented.

## Data availability

Single-cell RNA sequencing data have been uploaded to the Gene Expression Omnibus (Accession code GSE303380). To access for review, go to: https://urldefense.com/v3/ https://www.ncbi.nlm.nih.gov/geo/query/acc.cgi?acc=GSE303380;!!NZvER7FxgEiBAiR_!pAx7PWeubbltQaJ3VPlLelw-BI9-5SD65zJq78iUnb2JovhgDw_lAWtcj_iuPc6j8az5NbSTZqEw61lAQpGojR5XkaxrgDZ3$. Enter token **yrojuaccfjendkb** into the box. Code used for single-cell analysis is available at: https://github.com/Koehler-Lab/Amnoids-and-ventral-SkO-Study. Source data behind **Figures 1-3, 5, and Extended Data Figures 3-10** are available within the manuscript files.

## Supporting information

SIGuide

Source Data Ext-Data-Fig10

Source Data Ext-Data-Fig8

Source Data Ext-Data-Fig5

Source Data Ext-Data-Fig6

Source Data Ext-Data-Fig9

Source Data Ext-Data-Fig7

Source Data Ext-Data-Fig3

## Acknowledgments

K.R.K. is supported by National Institute of Health grants, R01AR075018 and R01DC017461. O.P. is supported by NIH RO1 grants R01HD113792 and 5R01HD085121-10. Fetal tissue specimens used in the Koehler Lab at Boston Children’s Hospital were obtained from the University of Washington Birth Defects Research Laboratory, which is supported by NIH award number 5R24HD000836 from the Eunice Kennedy Shriver National Institute of Child Health and Human Development, to I.A.G. We thank Muzlifah Haniffa and her team for assisting with transferring and annotation of human fetal skin single-cell sequencing atlas and for feedback on the project. We thank Clifford J Tabin for insights on the lateral plate mesoderm differentiation method. We thank Kimberly A. Aldinger, Dan Doherty, Ian G. Phelps, Jennifer C. Dempsey, Kevin J. Lee, Lucinda A. Cort for assistance in procuring fetal samples. We thank all the members of the Koehler Lab for their critical insights while preparing this manuscript.

## Author Contributions

A.P.L. and K.R.K. conceived and designed the study. A.P.L. conducted experiments, supervised experimental execution, wrote the initial manuscript draft, and edited the manuscript. J.K. designed and fabricated the microfluidic system, contributed to the initial manuscript draft, and edited the manuscript. Q.M. and L.S. performed single-cell RNA-seq data analysis and interpretation. K.Y.G., S.A.S., E.H.L., S.S, and Y.M. performed organoid differentiation, histology, and whole-mount staining. A.P.L., T.N., K.R.K., and O.P. developed the LPM differentiation approach. L.C.N., C.T.M., and B.I.L. assisted with procurement and processing of human skin specimens; B.I.L. additionally contributed to funding acquisition. I.A.G. provided human fetal tissue specimens. J.L. contributed to initial manuscript writing, editing, supervision, and funding acquisition. K.R.K. supervised the study, edited the manuscript, and secured funding. All authors reviewed and approved the final manuscript.

## Ethics Declarations

K.R.K. and J.L. are inventors of a patent relating to the skin organoid technology discussed in this article (WO2017070506A1). A.P.L. and K.R.K., with Children’s Hospital Corporation, have applied for a patent relating to Amnioid technology (PCT/US2024/033843; Priority Date 6.13.2023). K.R.K. is a consultant to STEMCELL Technologies, which has licensed the skin organoid technology.

## Supplementary Videos

**Supplementary Video 1:** Confocal live imaging from day 7 to day 12 showing the fusion of cSkO (*TJP1⁺*, green), derived from *TJP1-mEGFP* cells, with LPM (HAND1+, magenta), derived from *HAND1-mScarlet-I/TJP1-mEGFP* cells, to form cvSkO.

**Supplementary Video 2:** Confocal live imaging from day 7 to day 15 showing the fusion of cSkO, generated from *HAND1-mScarlet-I/TJP1-mEGFP* cells, with LPM, generated from *TJP1-mEGFP* cells, to form cvSkO. Note the progressive reduction of *Hand1⁺* cells in the cSkO as the organoid further differentiates, and the emergence of *TJP1⁺* cells from the LPM forming nascent vasculature that progressively envelops the assembloid.

**Supplementary Video 3:** Confocal live imaging of a cvSkO showing LPM-derived contributions (magenta, from *mTagRFP-T-TUBA1B*) to both the vasculature (marked by asterisks) and the dermal papilla (marked by arrowheads). The cvSkO was generated from Hand1-mScarlet-I/TJP1-mEGFP cells. Green indicates *TJP1⁺* epidermal cells. Magenta cytoplasmic signal represents *TUBA1B*+, while magenta nuclear signal reflects *HAND1* expression. Representative here showing cytoplasmic magenta signal, suggesting that the dermal papillae are from LPM - indicating the trunk skin identity.

**Supplementary Video 4:** 3D render of close-in images of vSkO’s hair follicle with measurements. Whole-mount staining with KRT5 (green) was performed to identify the hair follicles.

**Supplementary Video 5:** 3D render of close-in images of cSkO’s hair follicle with measurements. Whole-mount staining with KRT5 (green) was performed to identify the hair follicles.

**Supplementary Video 6**: Example of mechanical stress (Pa) differences between Amnioids and SE organoids (SEOs), illustrating the role of mesenchymal cells in buffering and maintaining low stress levels within the organoid to support tissue growth. Epithelial cells are labeled with TJP1-mEGFP. Live imaging was acquired at 30-minute intervals.

**Supplementary Video 7:** Mechanical measurements of epithelia (TJP1-mEGFP) and mesenchyme (mTagRFP-T-TUBA1B) in amnioids, including PIV-derived velocity fields, tension, and stress maps. The data reveal moderately coordinated movements and mechanical properties between the two layers, suggesting that the mesenchyme serves a dual role in buffering and transmitting mechanical signals during mechanotransduction.

**Supplementary Video 8:** Amnioid (day24-26 post-differentiation) undergoing mechanical collapse and recovery.

---

**Extended Data Figure 1.**
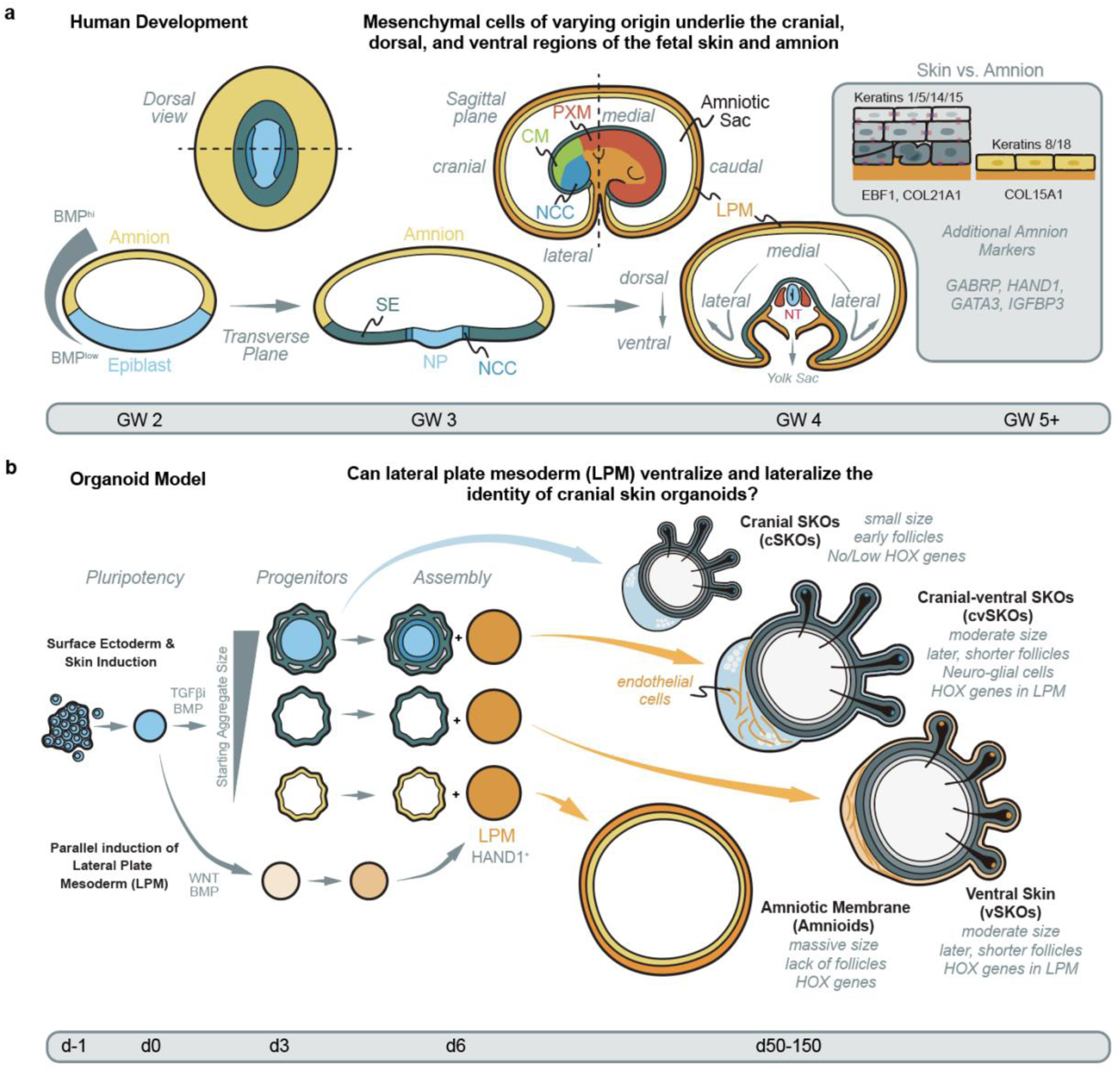
Developmental trajectories of human skin and amnion, and their recapitulation in organoid models. **a.** The establishment of regional skin during development. Mesenchymal cells of varying origins underlie the cranial, dorsal, and ventral regions of fetal skin and amnion. BMP signaling induces epiblast differentiation along the transverse plane, generating surface ectoderm (SE), neural plate (NP), and neural crest cells (NCC). Regional specification occurs through dorsoventral and mediolateral patterning, with distinct molecular markers distinguishing skin from amnion tissues (right inset). Paraxial mesoderm (PXM), cranial mesoderm (CM) **b.** Summary of *in vitro* model using pluripotent stem cells to generate organoids representing skin of specific anatomy. Parallel induction of surface ectoderm and LPM, followed by co-culture, generates cranial skin organoids with LPM derivatives (cvSkOs), ventral skin organoids (vSkOs) with moderate size, later-developing shorter follicles and *HOX gene* expression, and amniotic membrane models (Amnioids) characterized by absence of follicles.

**Extended Data Figure 2.**
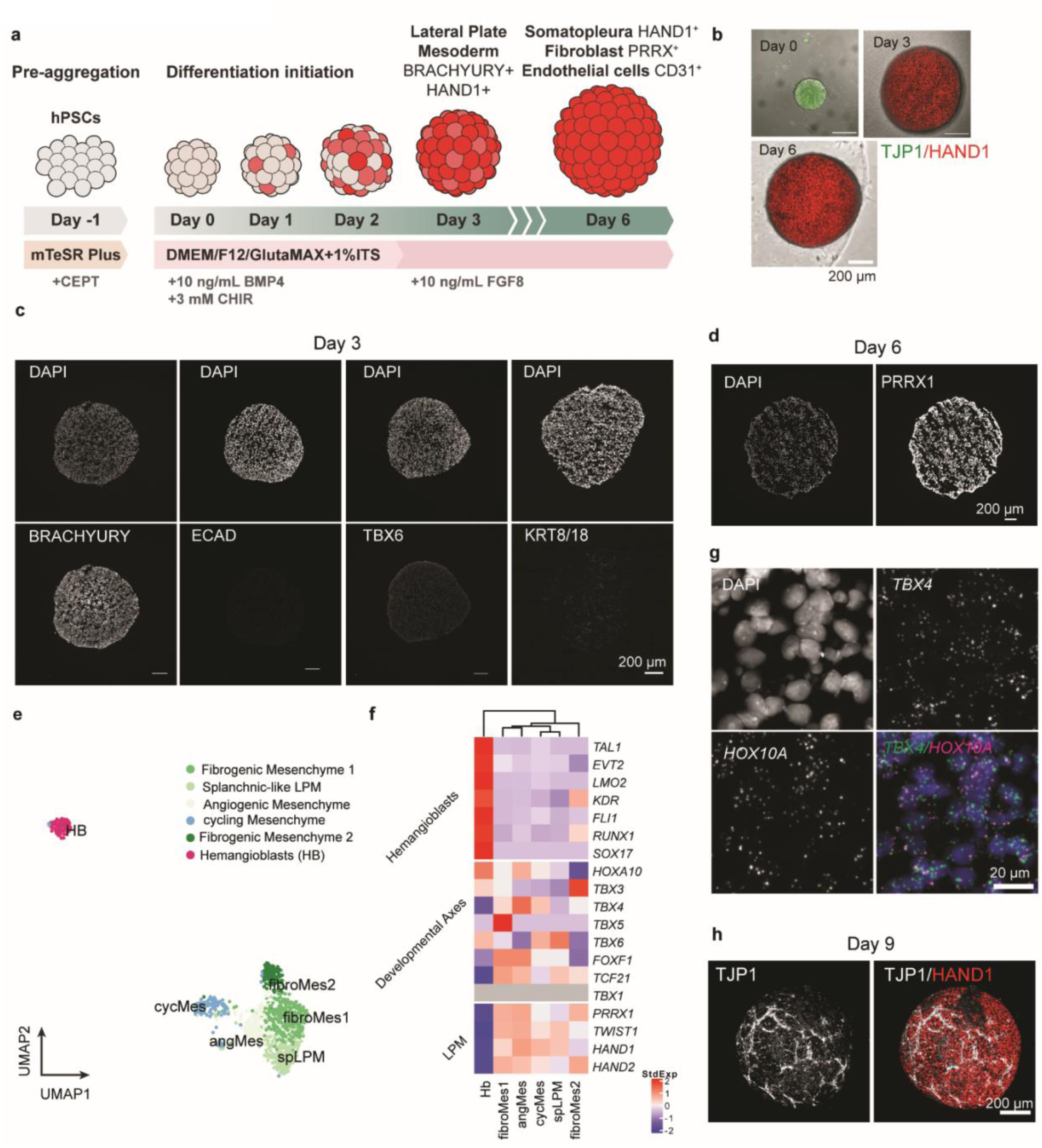
LPM Generation and characterization. **a.** Experimental workflow for generating lateral plate mesoderm organoids from human pluripotent stem cells. Pre-aggregated human pluripotent stem cell (hPSC) spheroids undergo differentiation from Day -1 to Day 6, with defined media conditions shown (mTeSR+, DMEMF12/GlutaMAX+ITS, supplemented with BMP4, CHIR, and FGF8). Progressive differentiation leads to LPM expressing BRACHYURY and HAND1 on day 3, and somatopleura fibroblasts (PRRX1+) and endothelial cells (CD31+) on day 6. **b.** Representative images of differentiating LPM spheroids at key timepoints (day 0, day 3, and day 6), showing morphological changes with HAND1 reporter (red) expression increasing over time. **c.** Immunofluorescence images of day 3 LPM showing expression of key mesodermal markers. **d.** Immunofluorescence images of day 6 spheroids showing PRRX1, a marker of lateral plate mesoderm derivatives. **e.** UMAP visualization of single-cell RNA sequencing of LPM at day 6, showing clusters of possible lineages emerged from LPM. **f.** Heatmap of selected markers indicating different lineages. *TBX1*, a marker of cardiopharyngeal mesoderm, was not expressed (gray). **g.** Fluorescent RNA *in situ* hybridization images of day 6 LPM showing regional marker representing posterior hindlimb and ventral fibroblasts (*TBX4*). Scale bar: 50 μm. **h.** Live confocal images showing endothelia emergence, with marked TJP1 expression in LPM at day 9 post-differentiation.

**Extended Data Figure 3.**
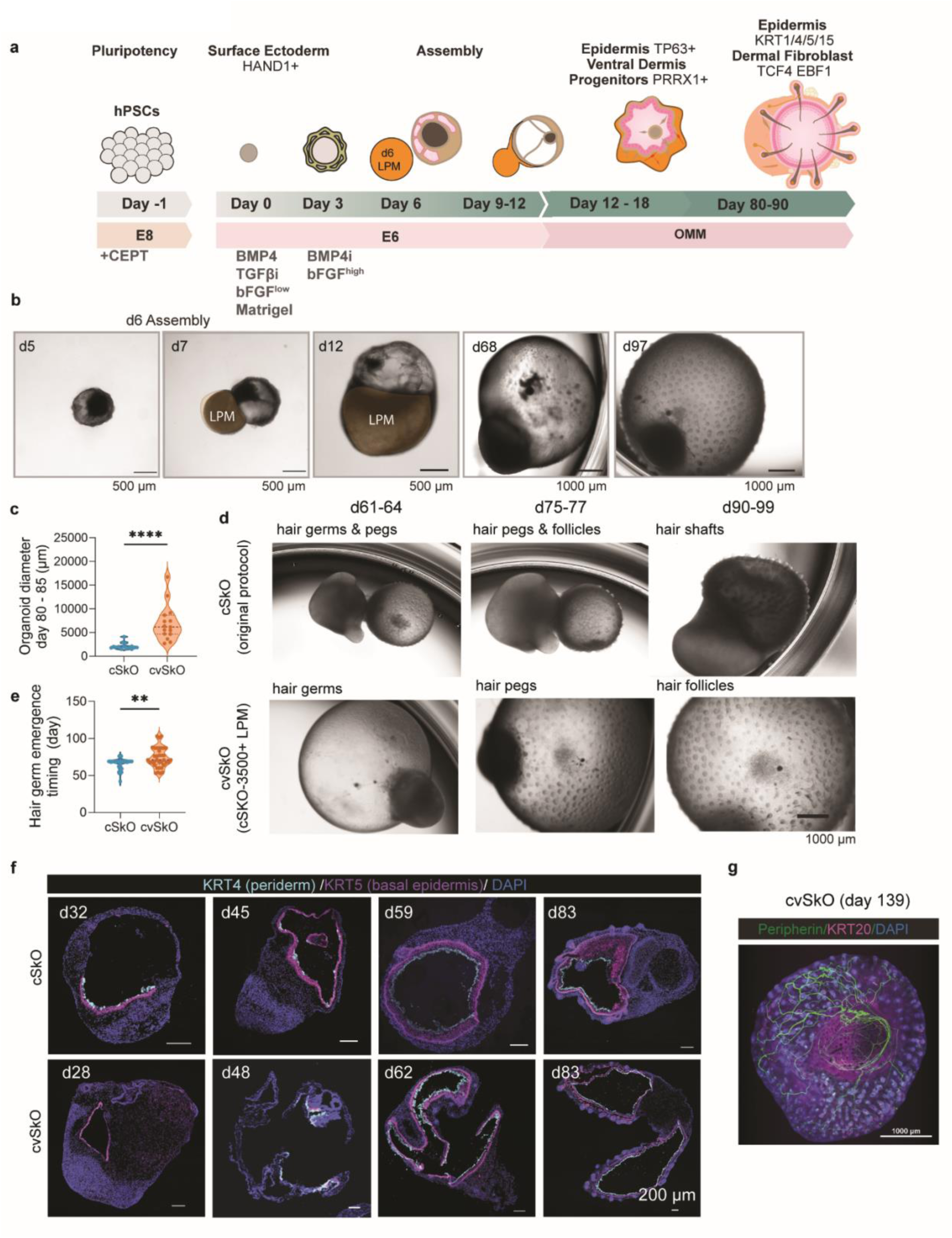
Generation and characterization of skin organoids with mixed cranial and ventral identities (cvSkOs). **a.** Schematic of skin organoids development from assembly with lateral plate mesoderm from hPSCs. These organoids have both cranial and ventral identities, and maturation into epidermis (TP63+/KRT14/15+) and ventral dermis progenitors (PRRX1+), ultimately developing into organoids with dermal fibroblasts (TCF4/EBF1+). **b.** Brightfield images showing temporal progression of organoid development. From d6 assembly with surface ectoderm and LPM (orange), through d12, d65, and d90 stages to d99 (Scale bars: 500 μm), where hair follicle structures are visible (last image, white arrow, scale bars: 50 μm). **c.** Size comparison of cSkO and cvSkO organoids. Upper panel: diameter measurements show significantly larger cSkO organoids. (n = 17 organoids from 3 independent batches). **d.** Comparison of hair follicle development in cSkO (cranial, top row) versus cvSkO (cranioventral, bottom row) organoids showing 10-14 days delay in hair follicle emergence timeline. **e.** Quantitative comparison of hair germ development (emergence date) in cSkO versus cvSkO. (cSkO: n = 21 organoids from 3 independent batches, cvSkO: n = 20 organoids from 3 independent batches). **f.** Immunofluorescence analysis of organoid cross-sections at progressive timepoints (d28-d83) comparing cSkO (top row) and cvSkO (bottom row). KRT4 (periderm, green), KRT5 (epidermis, red), and DAPI (nuclei, blue) staining delayed stratification and thickening of skin layers. Scale bars: 200 μm. **g.** Whole-mount staining of cvSkO at day 133 showing neuron (PRPH+) and Merkel cells (KRT20+), which are originated from cranial neural crest cells, showing that cvSkO has cranial identity.

**Extended Data Figure 4.**
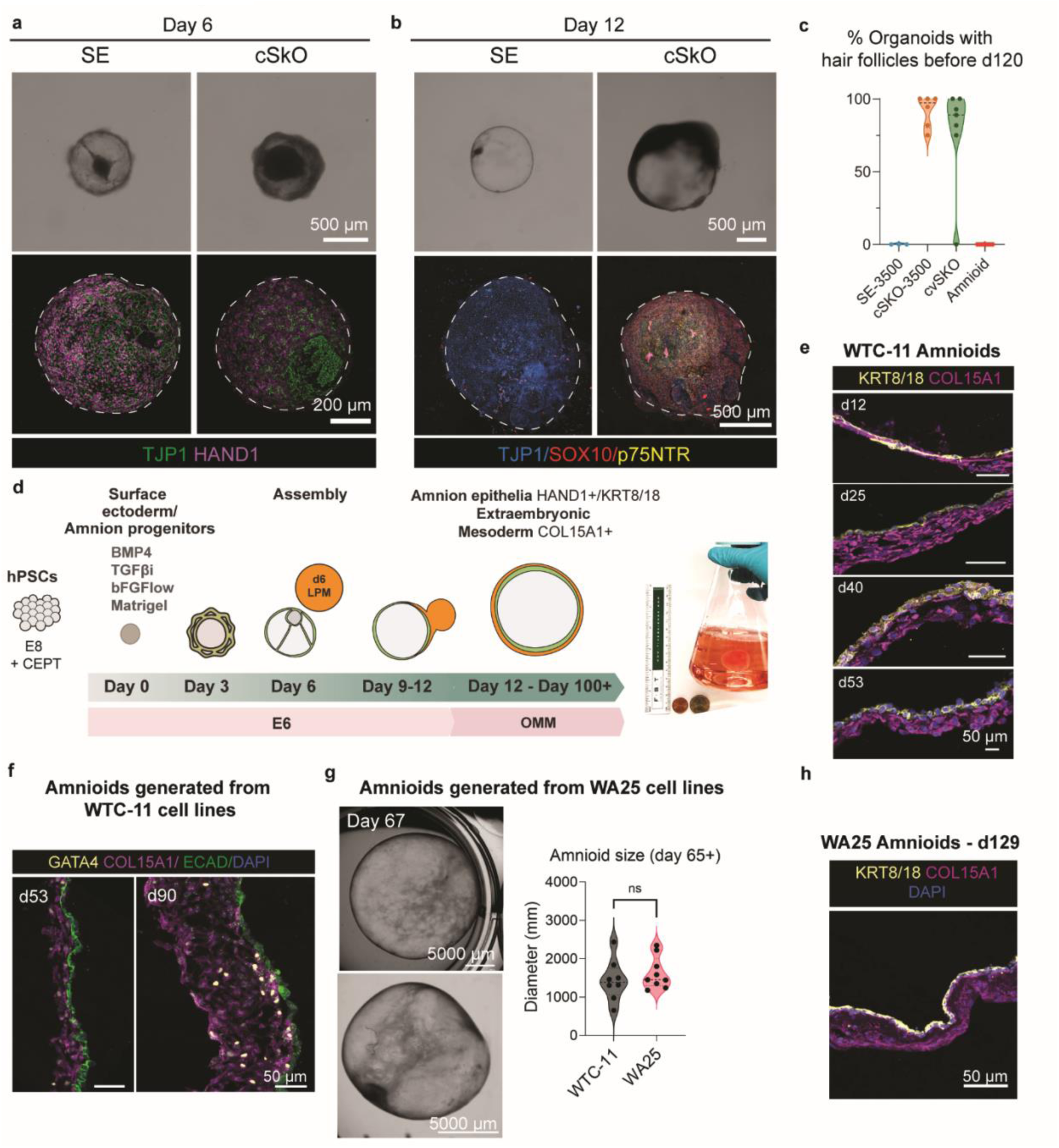
Characterization of Amnioids generated by modification to cvSkO protocol. **a.** Brightfield (top) and immunofluorescence (bottom) images of SEO and cSkO organoids at Day 6. TJP1 (green) and HAND1 (magenta) are shown. HAND1 expression is reduced in cSkO organoids, indicating impaired amnion differentiation. Dashed lines outline organoid boundaries. Scale bars: 200 μm. **b.** Brightfield (top) and immunofluorescence (bottom) images of SE and cSkO at Day 12 showing whole-mount staining for neural crest markers SOX10 (red) and p75NTR (yellow), along with ZO-1 (blue). Dashed lines outline organoid boundaries. Scale bars: 500 μm. **c.** Quantification of hair follicle formation. Violin plot shows the percentage of organoids forming hair follicles before Day 120 in SEOs, cSkOs, cSkOs, cvSkOs, and Amnioids. Folliculogenic capacity is restricted to SE-derived lineages. **d.** Schematic of the amnioid developmental protocol. Human pluripotent stem cells (hPSCs) are differentiated to amnion progenitors without neural crest induction, then assembled with Day 6 LPM to form amnioids. These undergo fusion, expansion, and mature into organoids (amnioids) comprising amnion epithelium (HAND1+, KRT8/18+) and extraembryonic mesoderm (COL15A1+). **Last panel:** Amnioids maintained in cornical flask at day 150+, reaching the size of about 3 cm diameter. **e.** Immunofluorescence time course of amnioid sections at Day 12, 25, 40, and 53, showing epithelial expansion (KRT8/18, white) and mesodermal marker COL15A1 (magenta). DAPI (blue) marks nuclei. **f.** Immunofluorescence of amnioids at Day 53 and Day 90 showing epithelial ECAD (green), extraembryonic mesoderm GATA4 (magenta), and DAPI (blue). **g.** Left: Examples of amnioids generated from WA25 cell line at Day 67, showing comparable size (quantified in Right panel) with amnioids generated from WTC cell lines. **h.** Immunofluorescence of amnioids generated from WA25 cell lines at Day 129 showing single epithelium (KRT8+) and COL15A1+ fibroblast layers.

**Extended Data Figure 5.**
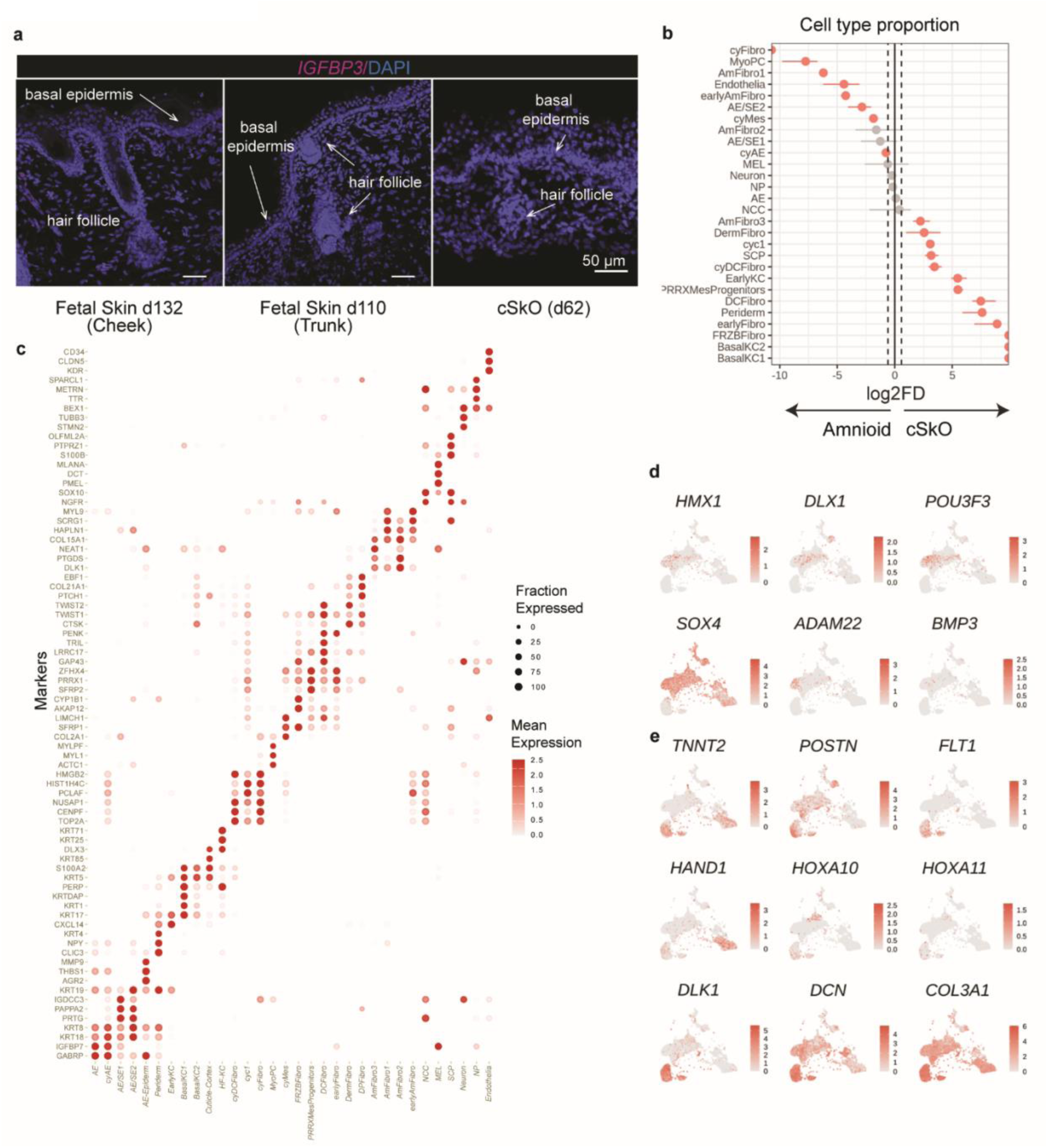
Single-cell transcriptomics comparison of Amnioids and cSkOs. **a.** *In-situ* hybridization (RNAScope) for *IGFBP3* in human fetal cheek skin (d132), trunk skin (d110), and cSkO (d62), showing absence of *IGFBP3* expression. **b.** Comparison of cell type proportions from day 28 single-cell transcriptomes of Amnioids and cSkOs. Amnioids are enriched in early mesenchymal and fibroblast populations, while cSkOs display increased neural crest and epidermal cell lineages. **b.** Cell type proportions comparison of single-cell transcriptomes of d28 Amnioid versus d28 cSkO. Amnioid models are enriched for early mesenchymal and fibroblast populations, while cSkOs show increased neural crest and epidermal lineage contributions. **c.** Dot plot showing expression of key marker genes across cell types identified in d28 Amnioids and cSkOs datasets. Dot size represents the fraction of cells expressing each marker; color intensity indicates mean expression level. **d.** Feature plots of genes enriched in cSkO fibroblasts based on DEG analysis, including cranio-pharyngeal markers (*HMX1, DLX1, POU3F3*), dermal fibroblast regulators (*SOX4*), and ECM modulators (*BMP3, ADAM22*), supporting cranial dermal identity and hair follicle-associated fate. **e.** Feature plots of genes enriched in Amnioid fibroblasts, including stress-response genes (*TNNT2, POSTN*), vasculature-associated marker (*FLT1*), mesodermal transcription factor (*HAND1*), ventral patterning genes (*HOXA10, HOXA11*), NOTCH regulator (*DLK1*), and extraembryonic ECM marker (*COL3A1*).

**Extended Data Figure 6.**
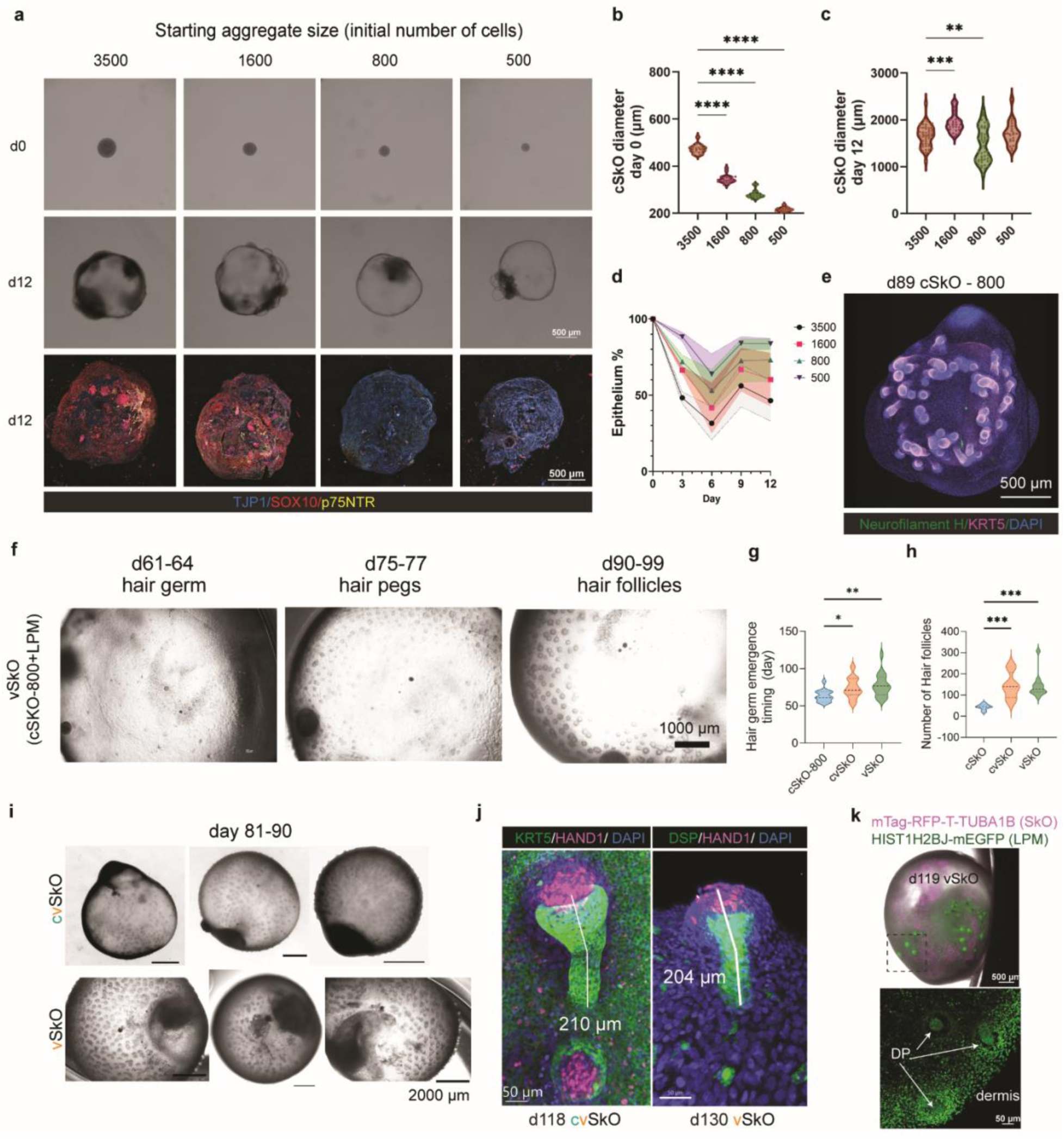
Reducing SkO starting cell size to generate purer ventral skin organoids. **a.** Brightfield (top and middle) and immunofluorescence (bottom) images of cSKO organoids derived from the WTC11 TJP1-mEGFP human iPSC line. Aggregates were generated using varying initial cell numbers (3,500; 1,600; 800; and 500 cells per aggregate) and imaged at day 0 and day 12 of differentiation. Immunostaining highlights epithelial (TJP1-mEGFP) and neural crest markers SOX10 and p75NTR. **b.** Quantification of cSkO diameter at day 0 across different starting cell numbers. (n = 20, 22, 22, and 19 organoids respectively for 3,500, 1,600, 800, and 500 cell conditions, from two independent experiments). **c.** Quantification of cSkO diameters at day 12 reveals no significant difference among the different starting cell numbers. **d.** Time-course analysis of epithelial area percentage from day 0 to day 12 shows differential epithelial expansion depending on initial aggregate size. **e.** Immunofluorescence of day 89 cSkO (800 cells as starting size), showing minimal neuronal contribution based on Neurofilament H staining. **f.** Brightfield images of vSkO organoids between days 61–99 illustrating sequential stages of hair follicle development (hair germ, hair peg, and mature follicle) with delayed emergence. **g.** Quantification of hair germ emergence timing in cSkO, cvSkO, and vSkO conditions; cSkO organoids exhibit earlier hair germ initiation (cSkO: n = 15 organoids in 3 batches, cvSkO: n = 36 organoids in 5 batches, vSkO = 17 organoids in 4 batches). Values used for cvSkOs are similar to those used in ExtDataFig.3e. **h.** Estimated number of observable hair follicles per organoid from brightfield images at day 80–90. LPM-infused organoids (cvSkO and vSkO) generate the highest number of follicles (cvSkO: n = 13 organoids in 5 batches, vSkO = 12 organoids in 4 batches). **i.** Brightfield images of vSkO and cvSkO organoids showing abundance of hair follicles. vSkO showing similar hair follicles emergence patterns as in cvSkO. Scale bar = 500 µm. **j.** Immunostaining of day 118 cvSkO and vSkO organoids showing hair follicle keratinocyte markers (KRT5 or DSP), dermal papilla marker HAND1 (LPM-derived), and DAPI, highlighting comparable hair follicle morphology between conditions. **k.** Whole-mount image of a day 119 vSkO organoid with RFP-labeled SkO and H2B-labeled LPM cells, indicating LPM contribution to both dermal fibroblasts and dermal papilla regions (white arrows). Insets showing zoom-in 20x images of hair follicles.

**Extended Data Figure 7.**
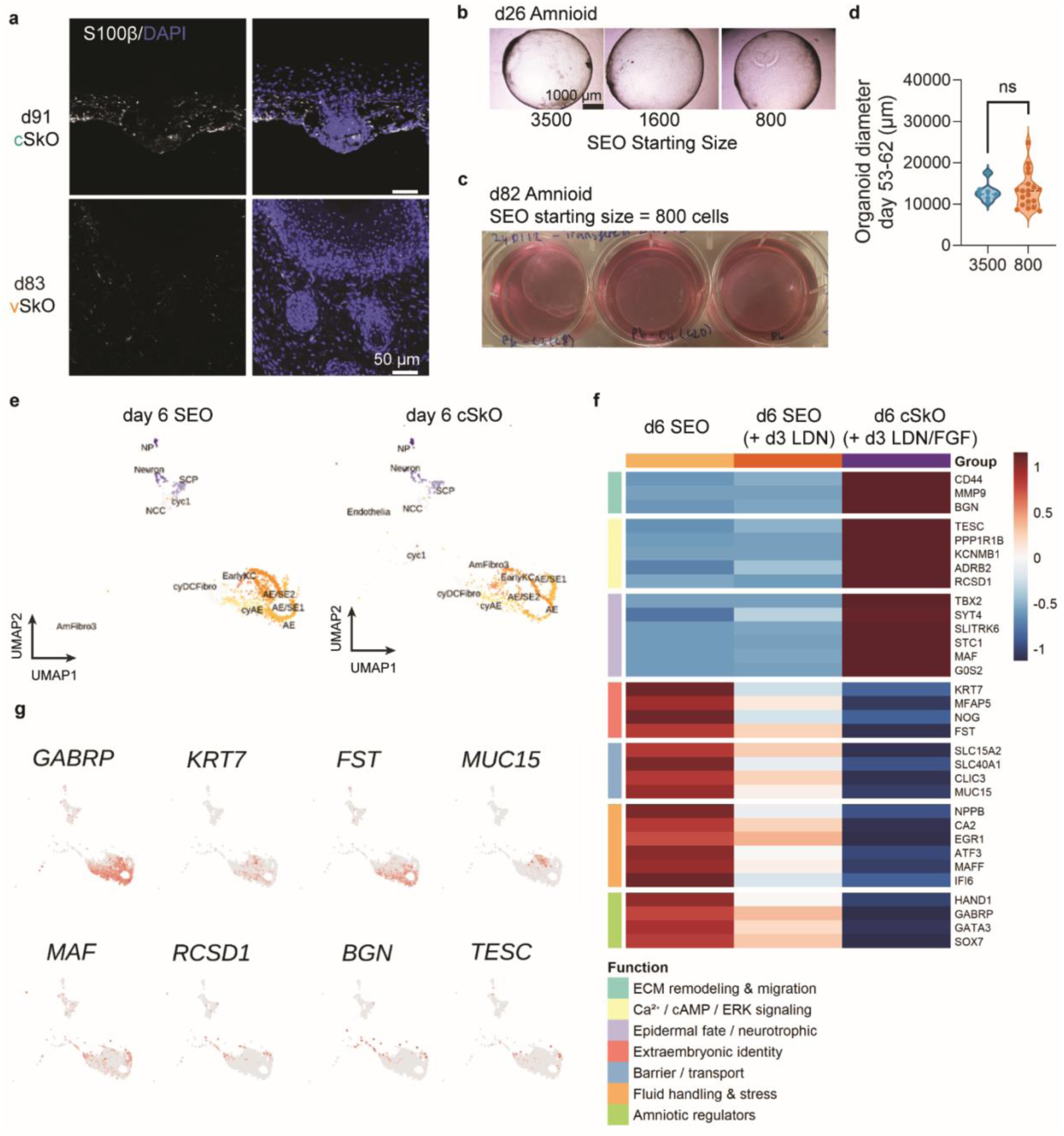
Schwann cell depletion, size scalability, and transcriptional divergence in early epithelial organoids. **a.** Immunofluorescence for S100β and DAPI in day 91 cSkO and day 83 vSkO reveals reduced Schwann cell presence in vSkO. Scale bars, 100 µm. **b.** Brightfield images of day 26 Amnioids seeded with 3500, 1600, or 800 cells per aggregate show comparable morphology across all starting cell numbers. **c.** Handphone images of day 82 Amnioids seeded at 800 cells reveal large organoid size similar to that of 3500-cell aggregates. **d.** Quantification of organoid diameter at day 53-62 showing similar amnioid size can be also be achieved using 800-cell starting aggregates (3500: n = 7 from 2 independent batches, 800: n = 19 from 3 independent batches). **e.** UMAP visualization of day 6 SEO and cSkO single-cell transcriptomes reveals divergence in epithelial and neural crest cell populations. **f.** Heatmap of differentially expressed genes (DEGs) comparing day 6 SEO (±LDN/FGF) and cSkO, grouped by functional gene categories. **g.** Feature plots of selected marker genes highlight transcriptional changes in amniotic epithelial and mesenchymal states following early patterning.

**Extended Data Figure 8.**
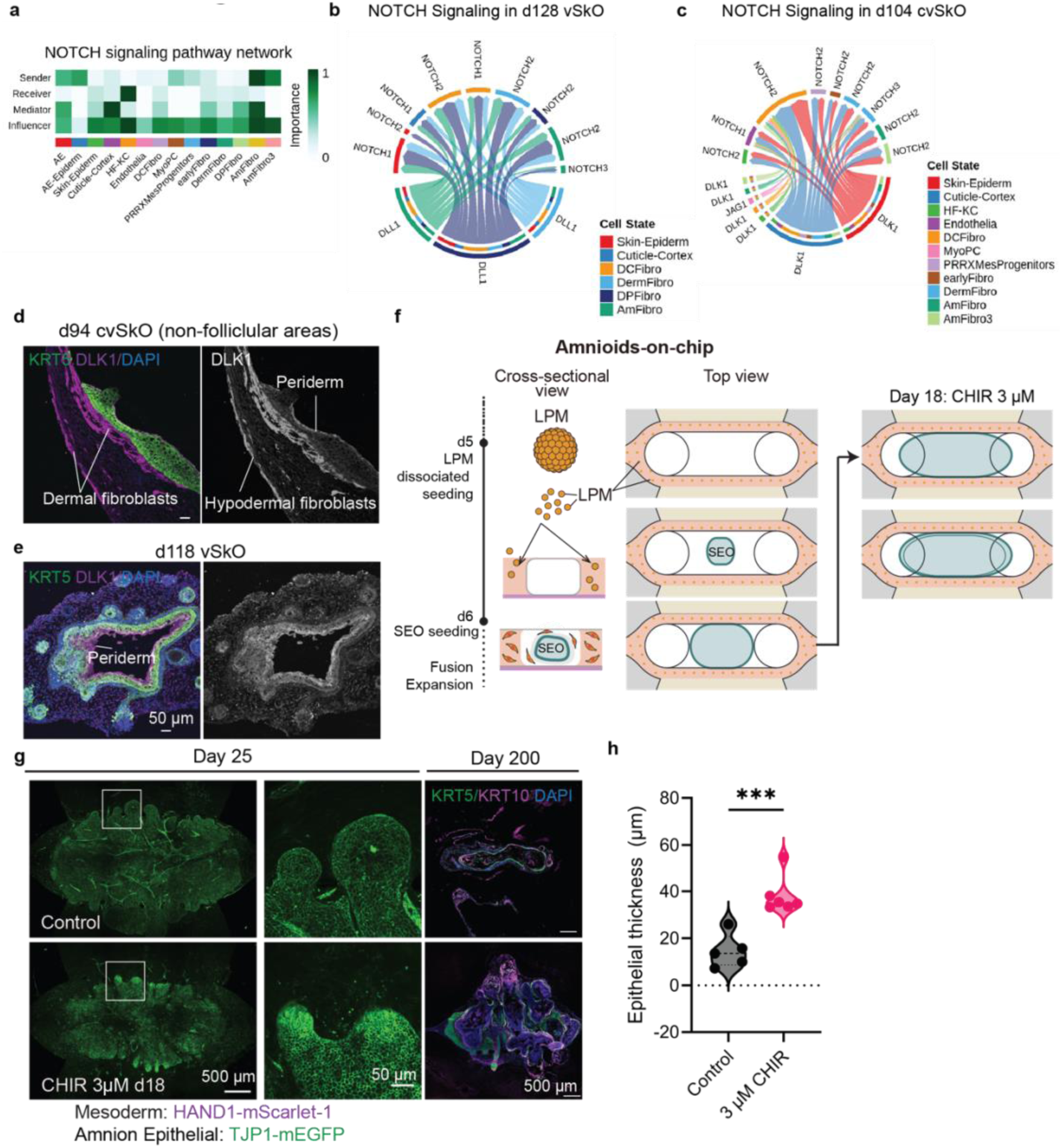
**a–c** CellChat analysis of NOTCH signaling pathway in day 100+ single-cell transcriptomic data. **a.** Network heatmap for NOTCH signaling showing overall signaling roles of cell clusters. **b.** Chord diagram comparing NOTCH ligands and receptors between vSkO and cvSkO; DLK1 emerges as a key driver of signaling differences. **c.** Chord diagrams comparing cvSkO and vSkO NOTCH signaling against that of cSkO, reveal cell-type specific pathway contributions. **d-e**. Immunofluorescence of (**d**) day 94 cvSkO (non-follicular region) and **(e)** day 118 vSkO stained for KRT5, DLK1, and DAPI. DLK1 marks dermal fibroblasts, hypodermal fibroblasts, and periderm. **f.** Schematic of amnioid-on-chip design showing sequential seeding of dissociated LPM at day 5 and SE epithelium at day 6, followed by 3 µM CHIR treatment on day 18 to induce amnion-to-skin transition. **g.** First three panels: wide-field (left) and zoom-in (right) views of day 25 amnioids-on-chip. CHIR treatment enhances epithelial stratification. HAND1-mScarlet marks mesoderm; TJP1-mEGFP marks epithelial cells. Scale bars, 500 µm (left), 50 µm (right). Last panel: immunofluorescent staining for sections from d200 chips, staining for KRT5 (basal) and KRT10 (intermediate) layers. CHIR-treated chip shows thickened keratinization. **h.** Quantification of epithelium thickness for day 25 chips shown in (f). n = 5 chips (Control) and n = 6 chips (CHIR-treated) in two independent experiments.

**Extended Data Fig. 9.**
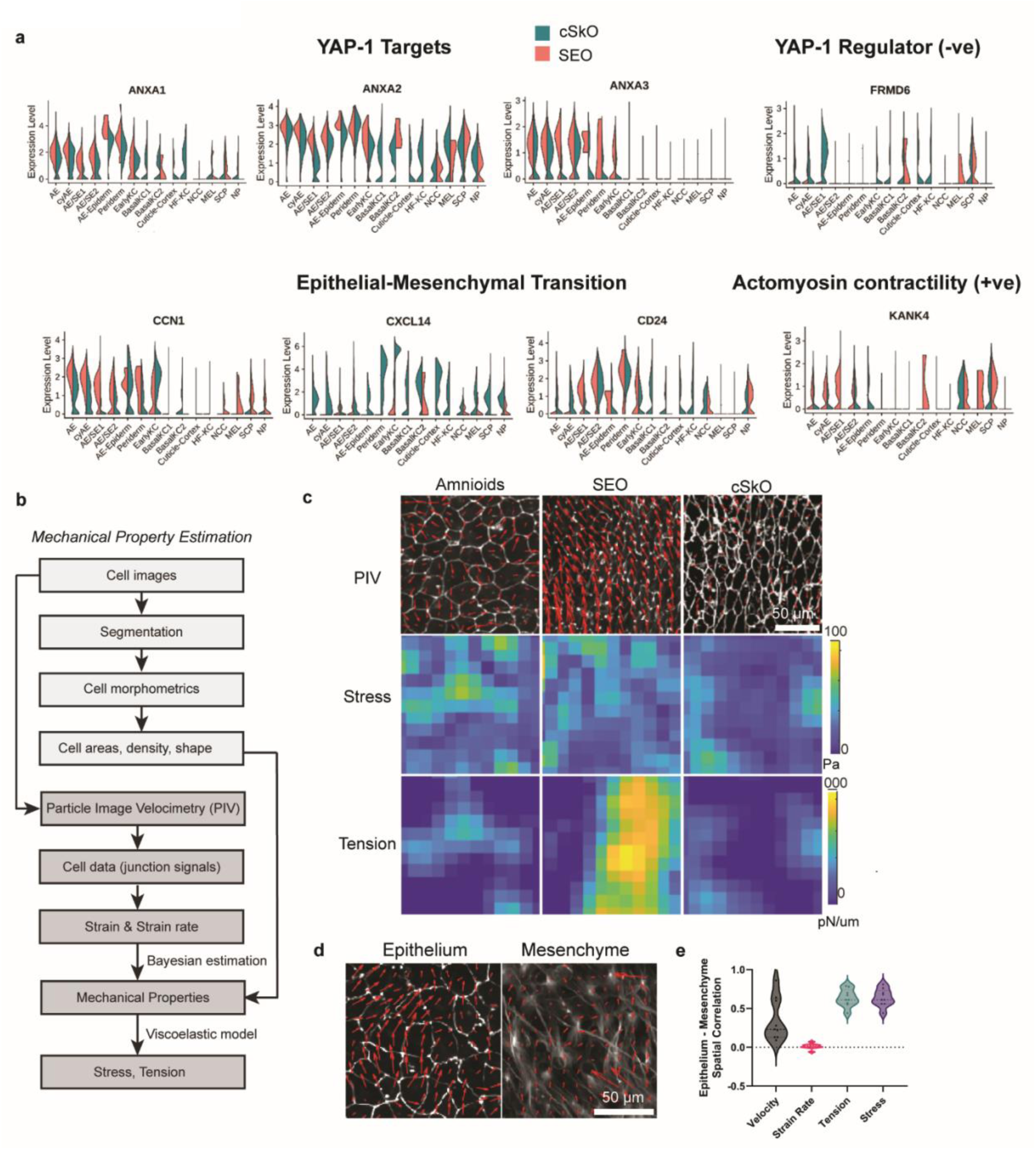
Molecular and mechanical features of epithelial–mesenchymal coupling in organoids. **a.** Violin plots showing expression of YAP1 targets (ANXA1–3), the YAP1 negative regulator FRMD6, epithelial–mesenchymal transition markers (CCN1, CXCL14, CD24), and actomyosin contractility gene KANK4 across multiple epithelial and mesenchymal populations. SEO (red) and cSKO (blue) groups are highlighted. (+ve): positive, (-ve): negative. **b.** Schematic workflow for mechanical property estimation from cell images using segmentation, morphometrics, Particle Image Velocimetry (PIV), and Bayesian modeling to compute stress and tension. **c.** Representative PIV displacement maps (top) and derived stress (middle) and tension (bottom) heatmaps in Amnioids, SEOs, and cSKOs. **d.** PIV velocity vector fields for epithelium and mesenchyme in Amnioids, showing moderate coupling of motion between layers. **e.** Quantification of epithelium–mesenchyme correlation for velocity and strain rate (low to moderate) versus stress and tension (high), suggesting mesenchyme functions both as a mechanical buffer and transmitter (n = 4 organoids in 2 independent batches).

**Extended Data Fig. 10.**
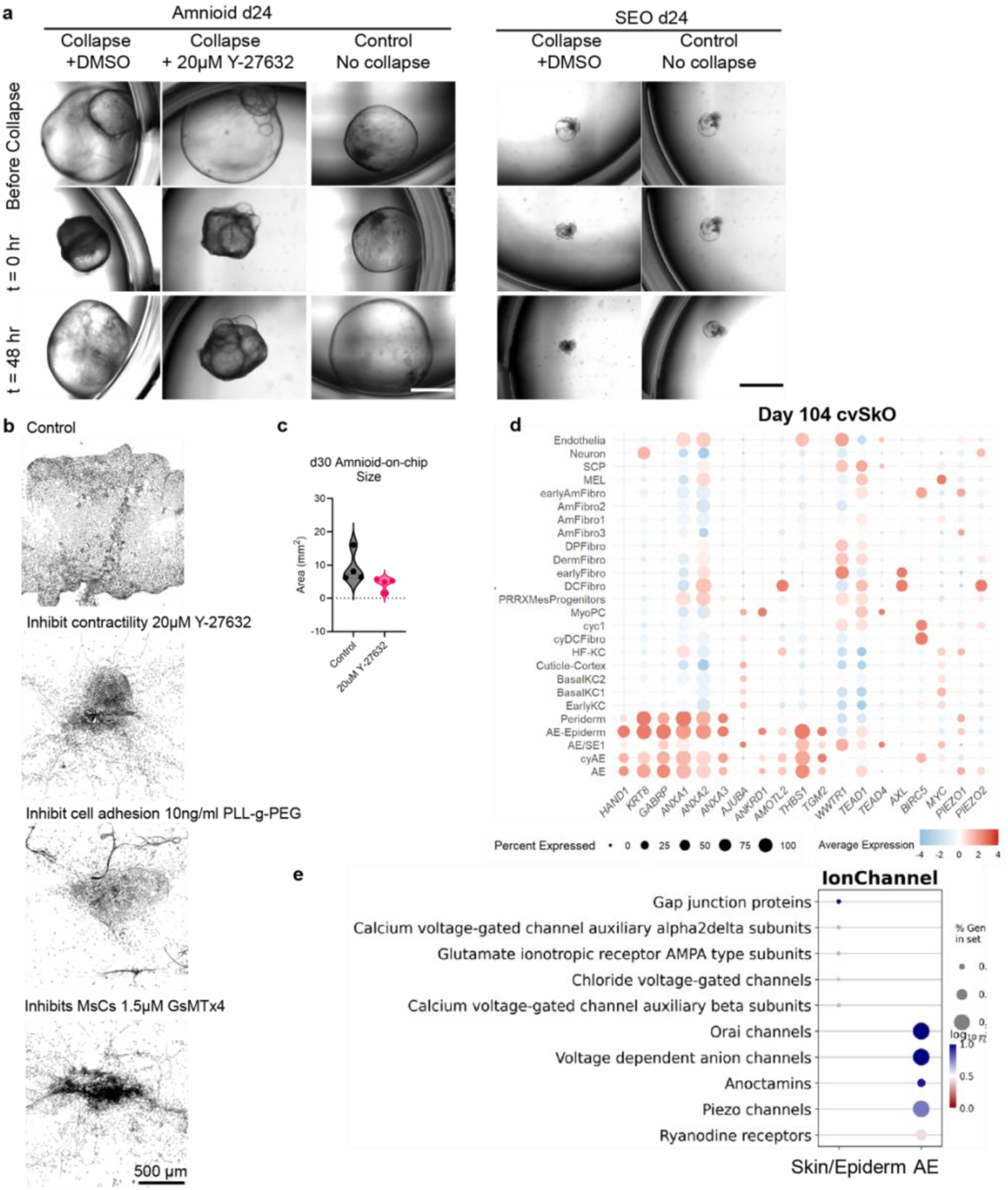
The role of LPM in amnioid recovery during mechanical perturbation and in cvSkOs’ mechanosensitive gene signatures. a. Brightfield images of Amnioids and SEs on day 24 under collapse-inducing conditions (±20 µM Y27632). ROCK inhibition prevents structural recovery in Amnioids. Images shown before collapse, immediately after (t = 0), and 48 hours post-collapse. **b.** Brightfield and ZO-1 confocal images of day 25 Amnioid-on-chip cultures treated with inhibitors. **c.** Quantification of Amnioid-on-chip area at day 30 comparing ROCK inhibitor and control conditions (n = 4 chips from 2 independent batches). **d.** Dot plots showing expression of amnion markers (*HAND1, KRT8, GABRP*), YAP targets, and mechanosensitive Piezo1/2 channels across cvSkO single-cell transcriptomes. Expression of YAP targets and Piezo channels is enriched in Amnion epithelia and transitional AE/SE clusters. The heatmap color scale reflects relative expression normalized to the dataset-wide average per gene. **e.** Gene set enrichment analysis (GSEA) of ion channel expression in amnion (AE, AE/SE1/2) versus epidermal clusters (earlyKC, Basal KC1/2, Periderm) across integrated single-cell datasets (day 6, 28, 100+). Amnion clusters show elevated expression of Piezo1/2 and calcium signaling channels (Orai, Ryanodine receptors).

## References

1. Jarzembowski, J. A. Pathobiology of Human Disease. Part II: Organ Syst. Pathophysiol.: Artic. Titles: N 2308–2321 (2014) doi:10.1016/b978-0-12-386456-7.05001-2.

2. Oldak, B. et al. Complete human day 14 post-implantation embryo models from naive ES cells. Nature 622, 562–573 (2023).

3. Aguilera-Castrejon, A. et al. Ex utero mouse embryogenesis from pre-gastrulation to late organogenesis. Nature 593, 119–124 (2021).

4. Weatherbee, B. A. T. et al. Pluripotent stem cell-derived model of the post-implantation human embryo. Nature 622, 584–593 (2023).

5. Rinn, J. L. et al. A dermal HOX transcriptional program regulates site-specific epidermal fate. Gene Dev 22, 303–307 (2008).

6. Driskell, R. R. et al. Distinct fibroblast lineages determine dermal architecture in skin development and repair. Nature 504, 277–281 (2013).

7. Minoux, M. & Rijli, F. M. Molecular mechanisms of cranial neural crest cell migration and patterning in craniofacial development. Development 137, 2605–2621 (2010).

8. Rinn, J. L. et al. A dermal HOX transcriptional program regulates site-specific epidermal fate. Genes Dev. 22, 303–307 (2008).

9. Rinn, J. L. et al. A Systems Biology Approach to Anatomic Diversity of Skin. J. Investig. Dermatol. 128, 776–782 (2008).

10. Rinn, J. L., Bondre, C., Gladstone, H. B., Brown, P. O. & Chang, H. Y. Anatomic Demarcation by Positional Variation in Fibroblast Gene Expression Programs. PLoS Genet. 2, e119 (2006).

11. Huang, D. et al. Lateral plate mesoderm cell-based organoid system for NK cell regeneration from human pluripotent stem cells. Cell Discov. 8, 121 (2022).

12. Tirosh-Finkel, L., Elhanany, H., Rinon, A. & Tzahor, E. Mesoderm progenitor cells of common origin contribute to the head musculature and the cardiac outflow tract. Development 133, 1943–1953 (2006).

13. Prummel, K. D., Nieuwenhuize, S. & Mosimann, C. The lateral plate mesoderm. Development 147, dev175059 (2020).

14. Glover, J. D. et al. The developmental basis of fingerprint pattern formation and variation. Cell 186, 940–956.e20 (2023).

15. Qi, L. et al. Krause corpuscles are genital vibrotactile sensors for sexual behaviours. Nature 630, 926–934 (2024).

16. Xiao, Z. et al. 3D reconstruction of a gastrulating human embryo. Cell 187, 2855–2874.e19 (2024).

17. Zhao, C. et al. A comprehensive human embryo reference tool using single-cell RNA-sequencing data. Nat. Methods 22, 193–206 (2025).

18. Tyser, R. C. V. et al. Single-cell transcriptomic characterization of a gastrulating human embryo. Nature 600, 285–289 (2021).

19. Shyer, A. E. et al. Emergent cellular self-organization and mechanosensation initiate follicle pattern in the avian skin. Science 357, 811–815 (2017).

20. Yang, S. et al. Morphogens enable interacting supracellular phases that generate organ architecture. Science 382, eadg5579 (2023).

21. Le, A. P., Kim, J. & Koehler, K. R. The mechanical forces that shape our senses. Development 149, (2022).

22. Aragona, M. et al. Mechanisms of stretch-mediated skin expansion at single-cell resolution. Nature 584, 268–273 (2020).

23. Jiang, C. et al. Mechanochemical control of epidermal stem cell divisions by B-plexins. Nat. Commun. 12, 1308 (2021).

24. Gopee, N. H. et al. A human prenatal skin cell atlas reveals immune cell regulation of skin morphogenesis. bioRxiv 2023.10.12.556307 (2023) doi:10.1101/2023.10.12.556307.

25. Lee, J. et al. Hair-bearing human skin generated entirely from pluripotent stem cells. Nature 582, 399–404 (2020).

26. Lee, J. et al. Generation and characterization of hair-bearing skin organoids from human pluripotent stem cells. Nat. Protoc. 17, 1266–1305 (2022).

27. Ramovs, V. et al. Characterization of the epidermal-dermal junction in hiPSC-derived skin organoids. Stem Cell Rep. 17, 1279–1288 (2022).

28. Basson, C. T. et al. Mutations in human cause limb and cardiac malformation in Holt-Oram syndrome. Nat. Genet. 15, 30–35 (1997).

29. Miao, Y. & Pourquié, O. Cellular and molecular control of vertebrate somitogenesis. Nat. Rev. Mol. Cell Biol. 25, 517–533 (2024).

30. Motazedian, A. et al. Multipotent RAG1+ progenitors emerge directly from haemogenic endothelium in human pluripotent stem cell-derived haematopoietic organoids. Nat. Cell Biol. 22, 60–73 (2020).

31. Hofbauer, P. et al. Cardioids reveal self-organizing principles of human cardiogenesis. Cell 184, 3299–3317.e22 (2021).

32. Prondzynski, M. et al. Efficient and reproducible generation of human iPSC-derived cardiomyocytes and cardiac organoids in stirred suspension systems. Nat. Commun. 15, 5929 (2024).

33. Holbrook, K. A. & Odland, G. F. Structure of the Human Fetal Hair Canal and Initial Hair Eruption. J. Investig. Dermatol. 71, 385–390 (1978).

34. Sudderick, Z. R., Glover, J. D., Batho-Samblas, C., Shih, B. B.-J. & Headon, D. J. Characterisation of human hair follicle development. bioRxiv 2024.07.26.605346 (2024) doi:10.1101/2024.07.26.605346.

35. Shao, Y. et al. Self-organized amniogenesis by human pluripotent stem cells in a biomimetic implantation-like niche. Nat Mater 16, 419–425 (2017).

36. Weatherbee, B. A. T. et al. Pluripotent stem cell-derived model of the post-implantation human embryo. Nature 622, 584–593 (2023).

37. Rostovskaya, M., Andrews, S., Reik, W. & Rugg-Gunn, P. J. Amniogenesis occurs in two independent waves in primates. Cell Stem Cell 29, 744–759.e6 (2022).

38. Pham, T. X. A. et al. Modeling human extraembryonic mesoderm cells using naive pluripotent stem cells. Cell Stem Cell 29, 1346–1365.e10 (2022).

39. Nguyen, H. et al. Tcf3 and Tcf4 are essential for long-term homeostasis of skin epithelia. Nat. Genet. 41, 1068–1075 (2009).

40. Miao, Q. et al. SOX11 and SOX4 drive the reactivation of an embryonic gene program during murine wound repair. Nat. Commun. 10, 4042 (2019).

41. Yu, Q. & Stamenkovic, I. Cell surface-localized matrix metalloproteinase-9 proteolytically activates TGF-β and promotes tumor invasion and angiogenesis. Genes Dev. 14, 163–176 (2000).

42. Zhang, H. et al. cAMP-PKA/EPAC signaling and cancer: the interplay in tumor microenvironment. J. Hematol. Oncol. 17, 5 (2024).

43. Kolobynina, K. G., Solovyova, V. V., Levay, K., Rizvanov, A. A. & Slepak, V. Z. Emerging roles of the single EF-hand Ca2+ sensor tescalcin in the regulation of gene expression, cell growth and differentiation. J. Cell Sci. 129, 3533–3540 (2016).

44. Gibson-Brown, J. J., Agulnik, S. I., Silver, L. M., Niswander, L. & Papaioannou, V. E. Involvement of T-box genes Tbx2-Tbx5 in vertebrate limb specification and development. Development 125, 2499–2509 (1998).

45. Reich, S., Kayastha, P., Teegala, S. & Weinstein, D. C. Tbx2 mediates dorsal patterning and germ layer suppression through inhibition of BMP/GDF and Activin/Nodal signaling. BMC Mol. Cell Biol. 21, 39 (2020).

46. Pham, T. X. A. et al. Modeling human extraembryonic mesoderm cells using naive pluripotent stem cells. Cell Stem Cell 29, 1346–1365.e10 (2022).

47. Hu, W. et al. Atlas of amnion development during the first trimester of human pregnancy. Nat. Cell Biol. 27, 1175–1185 (2025).

48. Sawada, R. et al. Analysis of drug transporter expression in syncytiotrophoblast derived from human placental stem cells: Expression and function of efflux transporters. Placenta 165, 23–32 (2025).

49. Rostovskaya, M., Andrews, S., Reik, W. & Rugg-Gunn, P. J. Amniogenesis occurs in two independent waves in primates. Cell Stem Cell 29, 744–759.e6 (2022).

50. Baek, J. et al. Egr1 is a 3D matrix–specific mediator of mechanosensitive stem cell lineage commitment. Sci. Adv. 8, eabm4646 (2022).

51. Gaut, L. et al. EGR1 Regulates Transcription Downstream of Mechanical Signals during Tendon Formation and Healing. PLoS ONE 11, e0166237 (2016).

52. Gunne-Braden, A. et al. GATA3 Mediates a Fast, Irreversible Commitment to BMP4-Driven Differentiation in Human Embryonic Stem Cells. Cell Stem Cell 26, 693–706.e9 (2020).

53. Gharibi, B. et al. Post-gastrulation amnioids as a stem cell-derived model of human extra-embryonic development. Cell (2025) doi:10.1016/j.cell.2025.04.025.

54. Andl, T., Reddy, S. T., Gaddapara, T. & Millar, S. E. WNT Signals Are Required for the Initiation of Hair Follicle Development. Dev. Cell 2, 643–653 (2002).

55. Ito, M. et al. Wnt-dependent de novo hair follicle regeneration in adult mouse skin after wounding. Nature 447, 316–320 (2007).

56. Driskell, R. R. et al. Distinct fibroblast lineages determine dermal architecture in skin development and repair. Nature 504, 277–281 (2013).

57. Nueda, M.-L. et al. DLK proteins modulate NOTCH signaling to influence a brown or white 3T3-L1 adipocyte fate. Sci. Rep. 8, 16923 (2018).

58. Finn, J. et al. Dlk1-Mediated Temporal Regulation of Notch Signaling Is Required for Differentiation of Alveolar Type II to Type I Cells during Repair. Cell Rep. 26, 2942–2954.e5 (2019).

59. Blanpain, C., Lowry, W. E., Pasolli, H. A. & Fuchs, E. Canonical notch signaling functions as a commitment switch in the epidermal lineage. Genes Dev. 20, 3022–3035 (2006).

60. Nier, V. et al. Inference of Internal Stress in a Cell Monolayer. Biophys. J. 110, 1625–1635 (2016).

61. Ishihara, S. & Sugimura, K. Bayesian inference of force dynamics during morphogenesis. J. Theor. Biol. 313, 201–211 (2012).

62. Sugimura, K., Bellaïche, Y., Graner, F., Marcq, P. & Ishihara, S. Robustness of force and stress inference in an epithelial tissue. arXiv 2013, 2712–2715 (2014).

63. Kastan, N. et al. Small-molecule inhibition of Lats kinases may promote Yap-dependent proliferation in postmitotic mammalian tissues. Nat. Commun. 12, 3100 (2021).

64. Hosseini, M., Koehler, K. R. & Shafiee, A. Biofabrication of Human Skin with Its Appendages. Adv. Healthc. Mater. 11, 2201626 (2022).

65. Hirsch, T. et al. Regeneration of the entire human epidermis using transgenic stem cells. Nature 551, 327–332 (2017).

66. Berg, S. et al. ilastik: interactive machine learning for (bio)image analysis. Nat. Methods 16, 1226–1232 (2019).

67. Aigouy, B., Umetsu, D. & Eaton, S. Segmentation and Quantitative Analysis of Epithelial Tissues. *Methods Mol. Biol. (Clifton*, NJ) 1478, 227–239 (2016).

