## Supplementary material for "Lateral plate mesoderm directs human amnion and ventral skin organoid formation": SIGuide

**SI GUIDE**

**Supplementary Information PDF:** Contains three Supplementary Notes on 1) Extended discussion of LPM induction optimization, 2) Extended discussion on assembloid generation, and 3) High-resolution scanning electron microscopy of amnioids and comparison with literature. Supplementary Method Figure 1 on flow cytometry gating scheme.

**Supplementary Tables**

**Supplementary Table 1**: List of antibodies, culture reagents and RNAScope probes.

**Supplementary Table 2:** Overall cell cluster annotations for the day-6 LPM dataset (sheet #1) and day-28-100+ mixed datasets (sheet #2).

**Supplementary Table 3:** Top 100 markers for each cell type.

**Supplementary Table 4:** Top differentially 100 genes in the comparison of Amnioid fibroblasts (earlyAmFibro, AmFibro1, AmFibro2, AmFibro3) and cSkOs fibroblasts (DCFibro, FRZBFibro, PRRXMesProgenitors, DermFibro, earlyFibro) clusters.

**Supplementary Table 5**: Ligand-Receptor signaling strength of NOTCH signaling pathway comparisons between cvSkO vs cSkO, vSkO vs cSkO and vSkO vs cvSkO.

**Supplementary Videos**

**Supplementary Video 1:** Confocal live imaging from day 7 to day 12 showing the fusion of cSkO (*TJP1⁺*, green), derived from *TJP1-mEGFP* cells, with LPM (*Hand1⁺*, magenta), derived from *Hand1-mScarlet-I/TJP1-mEGFP* cells, to form cvSkO.

**Supplementary Video 8:** Amnioid (day24-26 post-differentiation) undergoing mechanical collapse and recovery.

**Source Data Files (.xlsx format):** Figures 1, 2, 3, 5, and Extended Data Figs. 3-10
