## Supplementary Information for "Lateral plate mesoderm directs human amnion and ventral skin organoid formation"

**SUPPLEMENTARY NOTE**

**Supplementary Note 1: Extended discussion of LPM Induction optimization.**
We developed a protocol for lateral plate mesoderm (LPM) induction using 3 μM CHIR-99021 and BMP4 stimulation initiated on day 0. Protocol optimization was initially performed using HAND1-TJP1 reporter cells, guided by a set of morphological and molecular criteria described below. Following optimization, the protocol was primarily applied to the male WTC-11 iPSC line, and differentiation efficiency was assessed based on organoid morphology and optical features observed via bright-field microscopy. Secondarily, we validated this protocol using the female human ESC WA25 line using identical conditions (**Extended Data Fig. 4g**). Minor adjustments—such as BMP4 concentration or initial cell seeding density—may be needed for adaptation to other iPSC/ESC lines.

<u>**Criteria to assess LPM quality:**</u>

1. Bright-field imaging: By day 3, aggregates should exhibit a round, smooth, and opaque morphology, which should be maintained through day 6. By this point, fibroblasts typically begin to secrete extracellular matrix (ECM) and migrate outward, and endothelial differentiation may begin, producing translucent, epithelium-like regions.
2. HAND1 expression: HAND1 expression should be clearly detectable by fluorescent imaging by day 3 and should persist in the lateral plate mesoderm domain through at least day 10.
3. Immunohistochemistry (IHC) confirmation: By day 3, aggregates should express BRACHYURY and be negative for markers of alternative fates: PAX3/PAX6 (paraxial mesoderm/neural ectoderm), SOX17 (endoderm), and KRT8 (surface ectoderm). See Extended Data Fig. 2.
4. Quantitative metrics: Optimization was guided by quantifying aggregate roundness and HAND1 expression intensity within HAND1$^{+}$/TJP1$^{+}$ cells using immunofluorescence microscopy. To ensure consistency and unbiased quantification methods, images are taken at the same microscopy settings (with same lamp power, exposure, objectives). Also, epithelia versus mesenchyme bright-field segmentation was done on the same ilastik training.

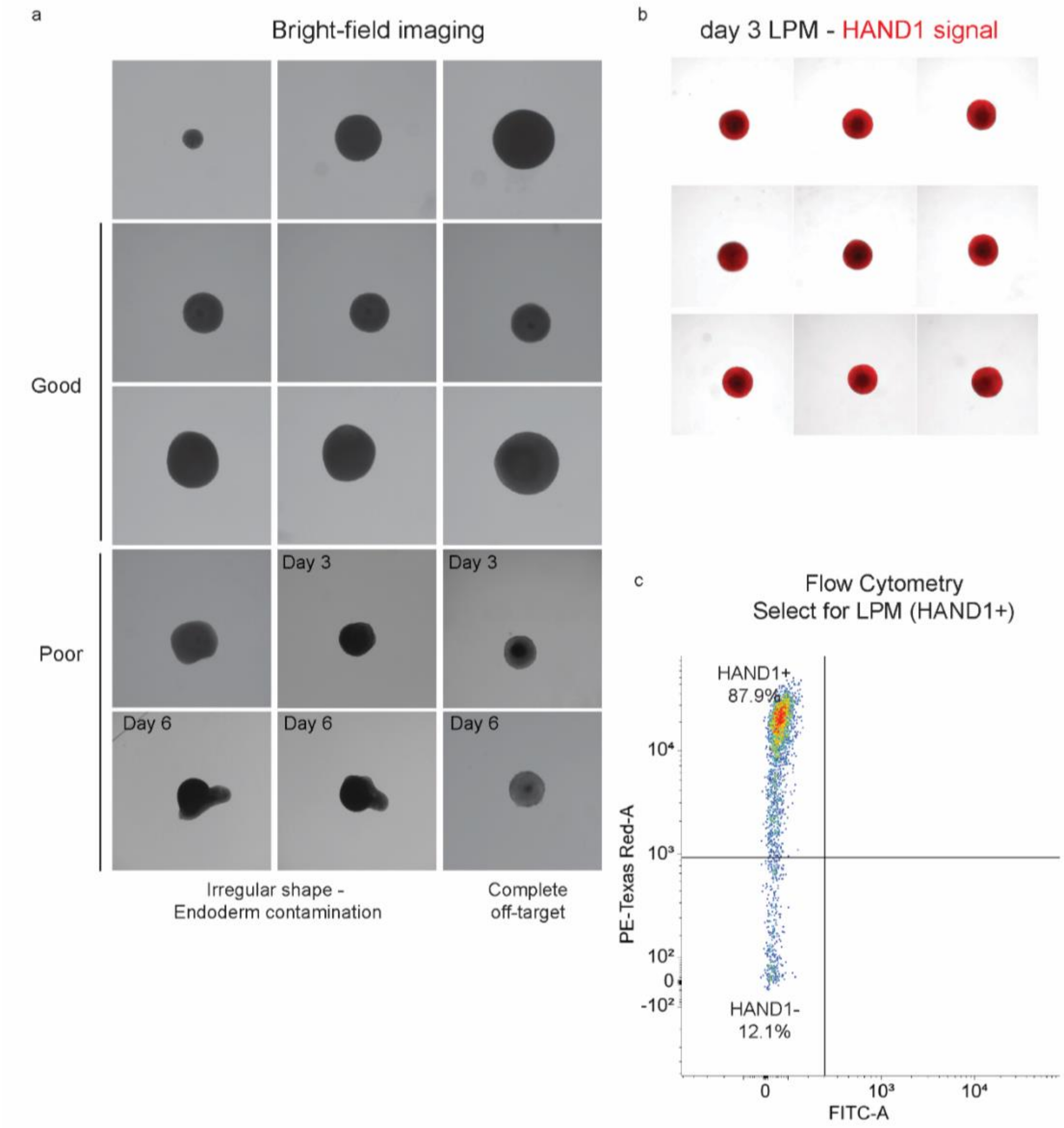


**Supplementary Fig.1: Example of visually identifiable efficient LPM production,** by HAND1+ signals and by bright-field imaging for first screening. Flow cytometry validation for HAND1+ (TexasRed channel) was performed to confirm.

**Serum conditions:** We tested serum added into the basal medium (DMEM/F12/GlutaMAX + 1% ITS) to promote endothelia and mesenchyme production. 0.1-5% Fetal Bovine Serum (FBS) or Knock-out serum replacement (KOSR) were tested.

- 0.5–1% FBS: Consistently supported smooth, round LPM structures and reproducible HAND1 expression.
- 5% FBS: Resulted in wrinkled, irregular morphologies indicative of endodermal bias.
- KnockOut Serum Replacement (KOSR): Tested as a serum alternative but showed inconsistent outcomes.

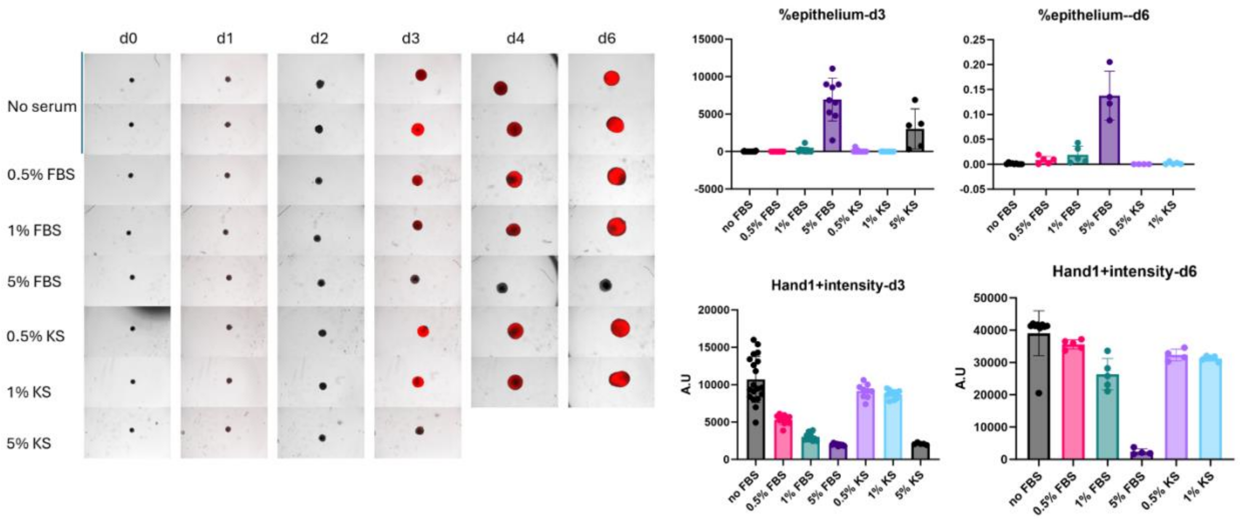


**Supplementary Fig. 2: Testing the addition of serum.**
To reduce variability, we ultimately adopted a serum-free protocol for LPM induction, which performed consistently across batches and was applied uniformly to all PSC lines.

**BMP4 concentration:** We compared 5 ng/mL and 10 ng/mL BMP4, applied on day 1 of differentiation. Both doses induced LPM features, but 10 ng/mL BMP4 yielded more consistent HAND1 expression and circular morphology. Thus, 10 ng/mL was selected for all subsequent LPM inductions.

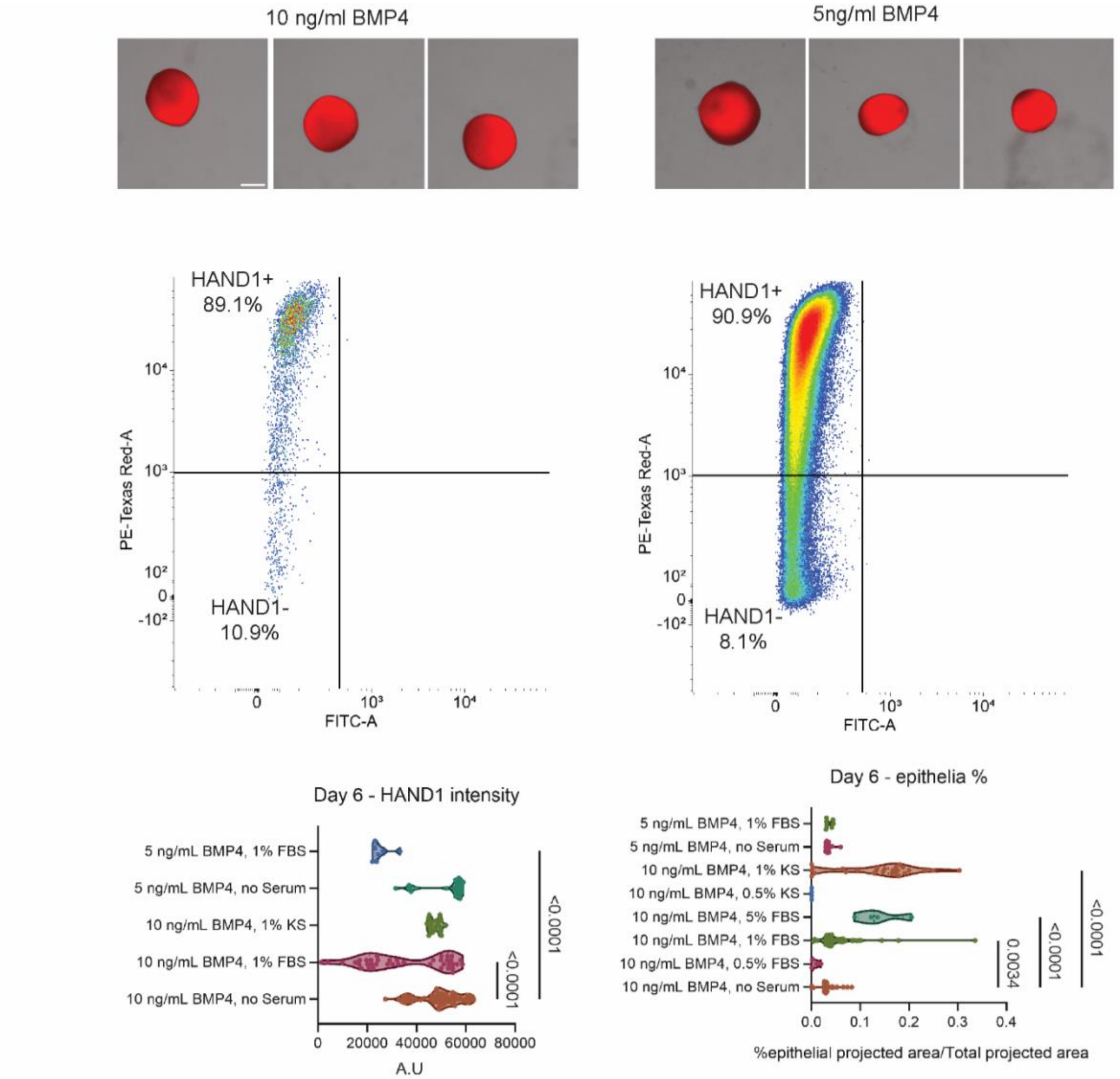


**Supplementary Fig.3: Testing BMP4 concentration.**

**<u>Starting cell density:</u>** We tested initial seeding densities of 800, 2,500, and 3,000 cells per aggregate. Starting cell number larger than 2500 cells per aggregate was capable of generating LPM-like aggregates; however, 3,000 cells per aggregate consistently produced the most optimal morphology and marker expression. In cases where a particular cell line exhibits a bias toward endoderm-like differentiation, increasing the initial cell number may help promote LPM identity. Conversely, if a radial gradient in ECAD/BRACHYURY expression is observed—suggesting cells at the edge of the aggregates are more exposed to BMP4 and adopting surface ectoderm-like fates—then reducing the starting cell number may help mitigate this effect. These observations underscore the importance of BMP4 diffusion and spatial signaling in aggregate patterning.

**Supplementary Note 2: Extended discussion of Assembloid Generation (Amnioids, cvSKO/vSKO lines)**

**LPM and SEO Combination to Generate Amnioids:**

**FBS vs. serum-free LPM:** When LPM generated with or without FBS was combined with SE to form Amnioids, both conditions resulted in equally efficient organoid formation and epithelial expansion.

**Mode of LPM incorporation:** We compared combination of cSkOs and SEO with whole LPM aggregates versus dissociated/sprinkled LPM cells during assembloid assembly. Both methods were equally effective in generating Amnioids. However, in vSKO and cvSKO lines, whole LPM aggregates more consistently yielded hair-bearing skin-like organoids, while dissociated LPM frequently gave rise to hairless, amnion-like epithelia.
To maximize reproducibility, we adopted the whole-organoid assembly approach as the standard.
**SEO generation:**

**LDN/FGF modulation:** We confirmed that FGF signaling was essential for SEO formation. Omission of FGF prevented SE identity. LDN was non-essential for SEO in our system and was excluded in the final protocol.

**Long-term Organoid Culture Medium:** We tested Advanced DMEM/F12 + GlutaMAX as a substitute for organoid maturation medium (OMM). This formulation supported long-term expansion and epithelial maintenance to day 30+ and was adopted as a cost-effective alternative without loss of efficiency.

**Supplementary Note 3: High-resolution scanning electron microscopy of Amnioids and comparison with literature.**

### Methods

Amnioids were carefully micro-dissected in standard OMM using fine forceps and spring scissors under a dissecting stereomicroscope (n=3 WTC-11 amnioids from one experimental batch). During the dissection, special attention was given to preserving the orientation of each cyst, distinguishing between the luminal surface (facing the internal cavity) and the exterior surface (facing the culture media). To immobilize the tissue, glass fibers previously adhered to 18 mm glass coverslips with Sylgard (Dow Corning) were used to gently pin the samples in place.

Tissues were fixed overnight at 4 °C in 4% paraformaldehyde (PFA) and then washed three times with ultrapure distilled water (Invitrogen) to remove residual fixative. Dehydration was performed through a graded ethanol series (30%, 50%, 70%, 95%, 100%), followed by a prolonged 48-hour incubation in 100% ethanol to ensure complete dehydration.

Samples were processed at the Harvard Electron Microscopy Core using an Autosamdri-931 critical point dryer (Tousimis) to preserve ultrastructural features. Dried samples were mounted on aluminum stubs using conductive carbon adhesive tabs and sputter-coated with 4 nm of platinum using a Leica EM ACE600 sputter coater. Scanning electron microscopy was performed using a Hitachi S-4700 field-emission scanning electron microscope (FE-SEM) at 5 keV acceleration voltage and 10 A extraction current. Images were acquired from both luminal and exterior surfaces of the organoids to capture regional surface differences and cellular morphology.

### Results

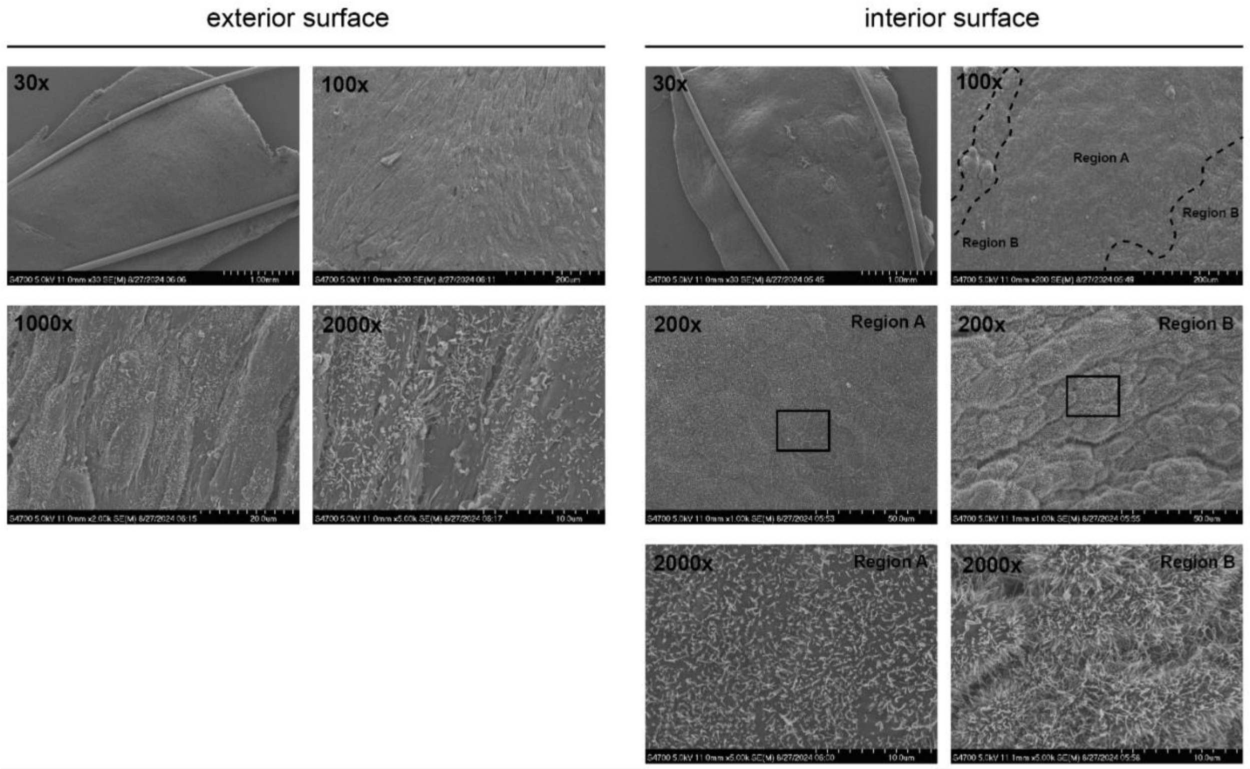

**Fig.4: Scanning Electron Microscopy of day 61 Amnioids** (generated from SEO-800 + LPM). Exterior surfaces show dense extracellular matrices (1000x) and cell debris (2000x), while interior surfaces (luminal) show organized epithelium layer with ordered cilia (2000x).

Scanning electron microscopy revealed distinct surface features between the interior (luminal) and exterior surfaces of the amnioids. The exterior surface exhibited irregular topography with accumulations of cell debris and dense extracellular matrix (ECM) components. These features were consistent with exposure to mechanical perturbation and the deposition of organic material during repeated media exchanges.

In contrast, the luminal surface displayed a more organized and consistent epithelial architecture. This surface was entirely ciliated, with regional differences in cilia length and density **(Fig.3, 2000x Region A & Region B).** Zones of tightly packed, shorter cilia interspersed with areas of longer, more sparsely spaced cilia. Underlying the cilia, the epithelial surface was composed of polygonal cells demarcated by cell–cell boundaries appearing slightly recessed relative to the apical surface.

Notably, we identified occasional large cells with expanded apical domains and prominent intracellular gaps crossed by thin cytoplasmic extensions **(Fig.3, 200x Region B)**. These features are characteristic of so-called "spider cells" previously described in human fetal amnion[1] . These cells appeared sporadically interspersed among smaller neighboring cells. In other regions, large, flat epithelial cells with shorter microvilli were observed in continuous sheets, with no evidence of intercellular canals.

### Discussion

Our scanning electron microscopy data provide compelling evidence that the in vitro amnioids recapitulate many key morphological features of the native human amnion, particularly those described in pre-term fetal samples. The presence of regional ciliation, polygonal epithelial organization, and specialized cell types such as spider cells mirrors descriptions from foundational studies of human amniotic epithelia[1,2]. This morphological fidelity supports the utility of these organoids as accurate structural models of the human amnion.

Previous ultrastructural analyses of diseased or abnormal amniotic tissues, including those from pregnancies complicated by polyhydramnios or omnihydramnios, have described notable disruptions to these characteristic features[2,3]. The ability to faithfully reproduce normal amniotic architecture in vitro now presents an opportunity to study such pathological deviations using patient-derived or genetically modified amniotic organoids.

Taken together, our findings highlight the translational potential of amniotic organoid systems for studying epithelial differentiation, barrier function, and aspects of amniotic disorders relevant to fetal development and maternal-fetal health. Future imaging studies of organoids modeling disease-associated conditions could provide insights into clinicopathologic features beyond those currently possible using human in vitro models.

**Supplementary Method Figure 1: Example of gating strategy for sorting LPM using cell sorter**

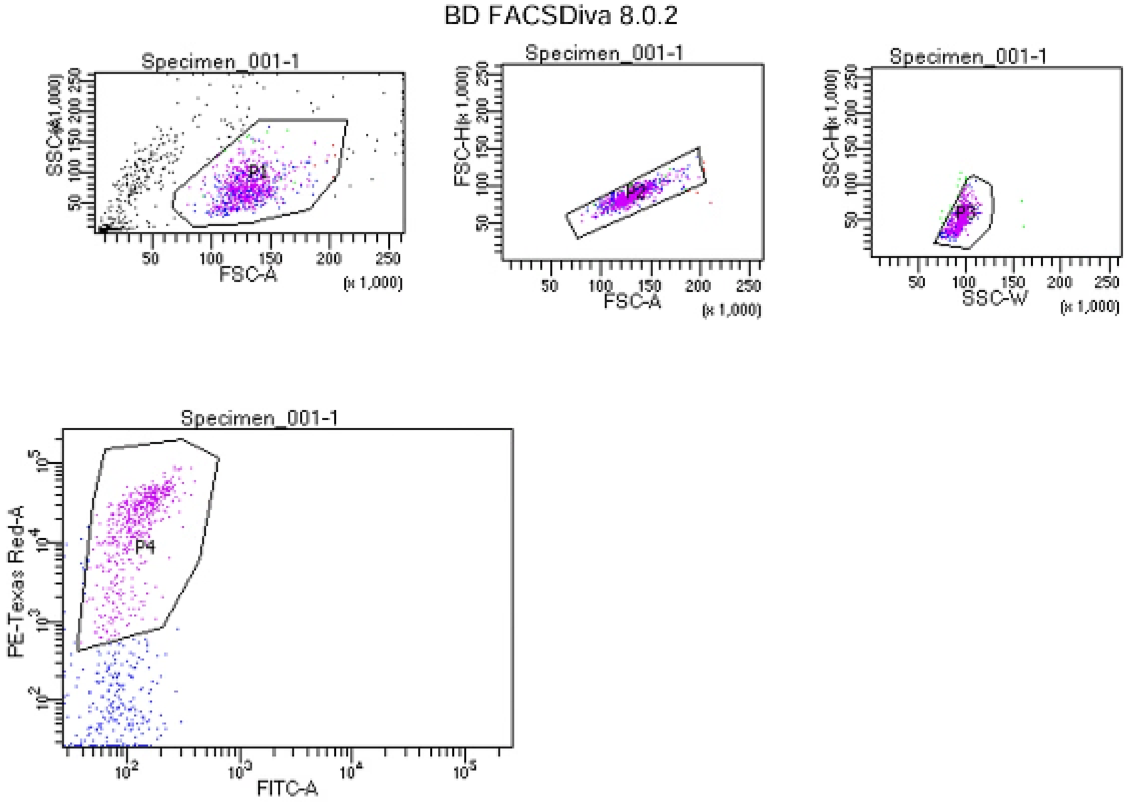


Tube: 1

| Population | #Events | %Parent | %Total |
|---|---|---|---|
| All Events | 1,316 | #### | 100.0 |
| P1 | 872 | 66.3 | 66.3 |
| P2 | 866 | 99.3 | 65.8 |
| P3 | 848 | 97.9 | 64.4 |
| P4 | 580 | 68.4 | 44.1 |

### Supplementary References

1. Pollard, S. M., Aye, N. N. & Symonds, E. M. Scanning electron microscope appearances of normal human amnion and umbilical cord at term. *Br. J. Obstet. Gynaecol.* **83**, 470–7 (1976).

2. Pollard, S. M., Symonds, E. M. & Aye, N. N. SCANNING ELECTRON MICROSCOPIC APPEARANCES IN THE AMNION IN POLYHYDRAMNIOS AND OLIGOHYDRAMNIOS. *BJOG: Int. J. Obstet. Gynaecol.* **86**, 228–232 (1979).

3. Hebertson, R. M., Hammond, M. E. & Bryson, M. J. Amniotic epithelial ultrastructure in normal, polyhydramnic, and oligohydramnic pregnancies. *Obstet. Gynecol.* **68**, 74–9 (1986).
